# How Many Streamlines are Required for Reliable Probabilistic Tractography? Solutions for Microstructural Measurements and Neurosurgical Planning

**DOI:** 10.1101/2020.01.26.920397

**Authors:** Lee B. Reid, Marcela I. Cespedes, Kerstin Pannek

## Abstract

Diffusion MRI tractography is commonly used to delineate white matter tracts. These delineations can be used for planning neurosurgery or for identifying regions of interest from which microstructural measurements can be taken. Probabilistic tractography produces different delineations each time it is run, potentially leading to microstructural measurements or anatomical delineations that are not reproducible. Generating a sufficiently large number of streamlines is required to avoid this scenario, but what constitutes “sufficient” is difficult to assess and so streamline counts are typically chosen in an arbitrary or qualitative manner. This work explores several factors influencing tractography reliability and details two methods for estimating this reliability. The first method automatically estimates the number of streamlines required to achieve reliable microstructural measurements, whilst the second estimates the number of streamlines required to achieve a reliable binarised trackmap than can be used clinically. Using these methods, we calculated the number of streamlines required to achieve a range of quantitative reproducibility criteria for three anatomical tracts in 40 Human Connectome Project datasets. Actual reproducibility was checked by repeatedly generating the tractograms with the calculated numbers of streamlines. We found that the required number of streamlines varied strongly by anatomical tract, image resolution, number of diffusion directions, the degree of reliability desired, the microstructural measurement of interest, and/or the specifics on how the tractogram was converted to a binary volume. The proposed methods consistently predicted streamline counts that achieved the target reproducibility. Implementations are made available to enable the scientific community to more-easily achieve reproducible tractography.

## 1 Introduction

Diffusion MRI measures the Brownian motion of water molecules in the brain, to which mathematical models can be applied to estimate the underlying orientation of white matter fibers. Tractography can then be applied to this model to delineate white matter pathways. Commonly, the scientific motivation for tractography is to sample microstructural measurements, such as fractional anisotropy (FA), of specific white matter tracts for the purpose of comparing populations or assessing changes over time (e.g. 1,2). An alternative motivation is to use the tractogram to guide neurosurgical planning or make morphological measurements (3,4). Probabilistic tractography is a popular means of performing these tasks but, unlike deterministic tractography, produces different delineations each time it is run. This presents an issue in both clinical and scientific contexts. If tractography is unreliable, microstructural measurements may be unreliable, potentially inflating Type I or Type II errors. More seriously, unreliable tractography in a clinical context might threaten patient safety (for example by underestimating the size of a tract, an issue also seen with deterministic tractography) or, at the very least, reduce the perceived usefulness of this tool for clinicians.

A major key to reliable probabilistic tractography is the number of streamlines generated. If two probabilistic tractograms are created with the same parameters, their streamline densities in corresponding voxels should converge as the number of streamlines increases. Extremely large numbers of streamlines, however, have high computational requirements to generate, view, and store. By contrast, whilst low streamline-count tractograms are less computationally expensive, even a cursory visual comparison against a higher streamline-count tractogram can demonstrate a failure to adequately delineate the desired anatomy (Figure 1). Some investigators have reported on the relationship between *whole-brain* streamline count and reproducibility in connectivity analyses (5–7), but for the anatomical delineation of specific tracts, little advice exists within the community for selecting a sensible number of streamlines. This is because the optimum number presumably relies on many factors, such as the head size, anatomy in question, sequence parameters, and image quality. Consequentially, the number of streamlines reported in published literature varies greatly, and authors rarely provide evidence that the streamline count chosen was sufficient to reliably delineate the anatomy in question.

**Figure 1.**
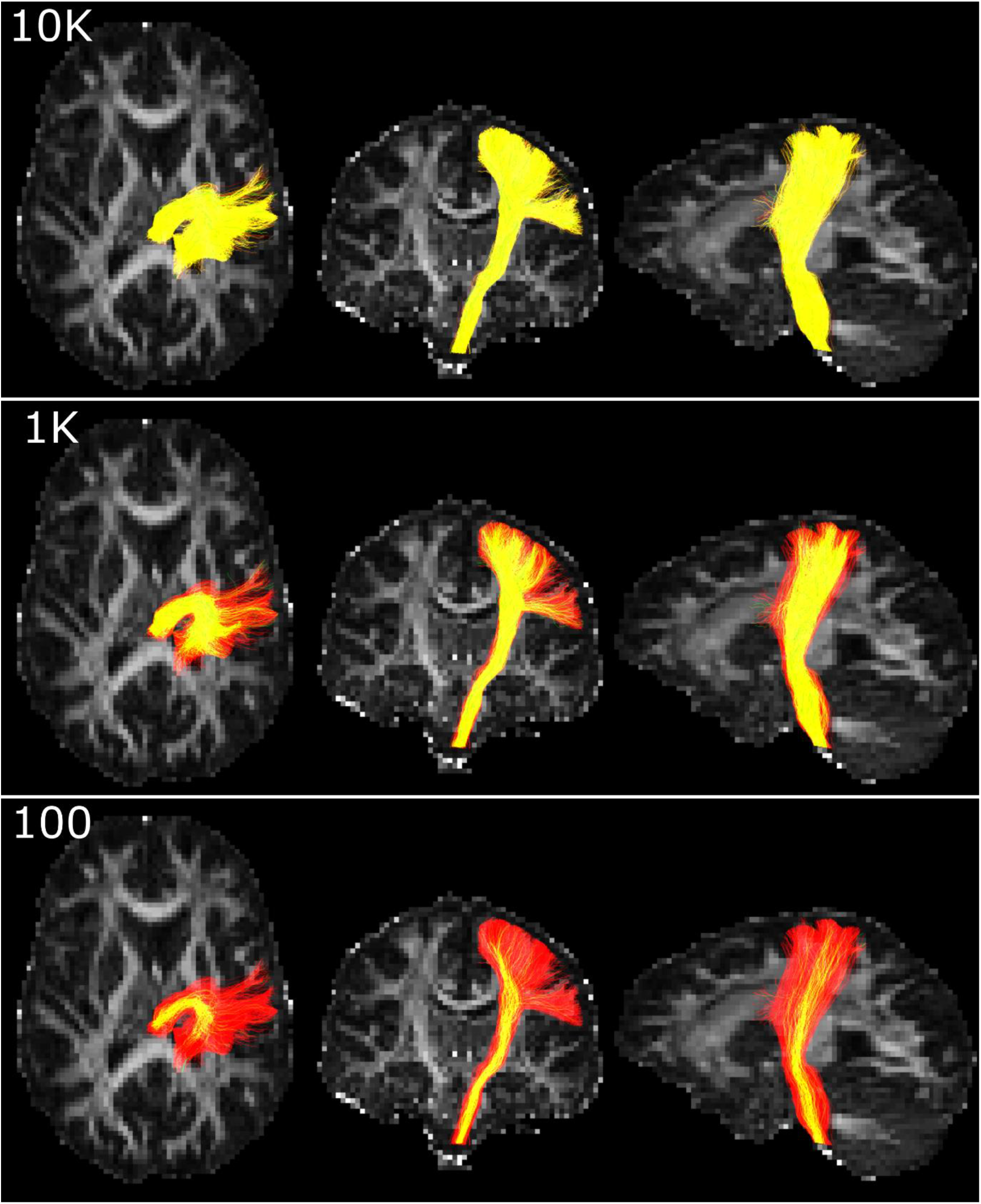
Tractograms of the corticospinal tract with 20,000 streamlines (red) overlaid with tractograms of 10,000 streamlines (top), 1000 streamlines (middle) and 100 streamlines (bottom). Yellow indicates overlap between the smaller and larger tractograms. The background image indicates fractional anisotropy. The 10,000 streamline tractogram predominantly overlaps the 20,000 streamline tractogram. By contrast, the smaller tractograms underestimate the extent of the corticospinal tract and suggest low confidence in its superior and anterior aspects that are reliably delineated by the larger tractogram. Data from Reid et al (12).

A potentially compounding issue is that, in a neurosurgical context, the ideal means of interpreting tractography is not necessarily in its raw form but may be as a binarised trackmap (a track-density image (8) that has been thresholded then binarised). Several arguments exist for the conversion of probabilistic tractography into this format. For example, overlaying images with raw streamline files or non-binary voxelwise representations thereof is not well supported by Picture Archiving and Communications System (PACS) oriented DICOM viewers that are central to clinical workflows (9,10). Clinicians are also generally more familiar with the more simplistic visualisations from deterministic diffusion tensor tractography that historically have dominated tractography research in surgical journals (for a review, see (11)). By contrast, viewing and interpreting probabilistic tractograms requires considerable experience, particularly with regards to judging the location of the true anatomical boundary, which can be obscured by false-positive streamlines, underestimated by obtaining too few streamlines, and modified by changing the depth of focus or transparency (Figures 1 & 2). Binary maps, by contrast, leave little room for misinterpretation. Most importantly, tissue resection is itself a binary operation. Surgeons need to make binary decisions and so it is appropriate that risk boundaries are delineated as such, particularly when an intuitive mathematical basis for formalizing such boundaries is available. Of course, the act of thresholding and binarising trackmaps can compound the difficulty in choosing an optimal number of streamlines. This is not only because a range of thresholding rules can be used but more importantly, to the best of our knowledge, the impact of such thresholding on reproducibility has yet to be formally documented.

**Figure 2.**
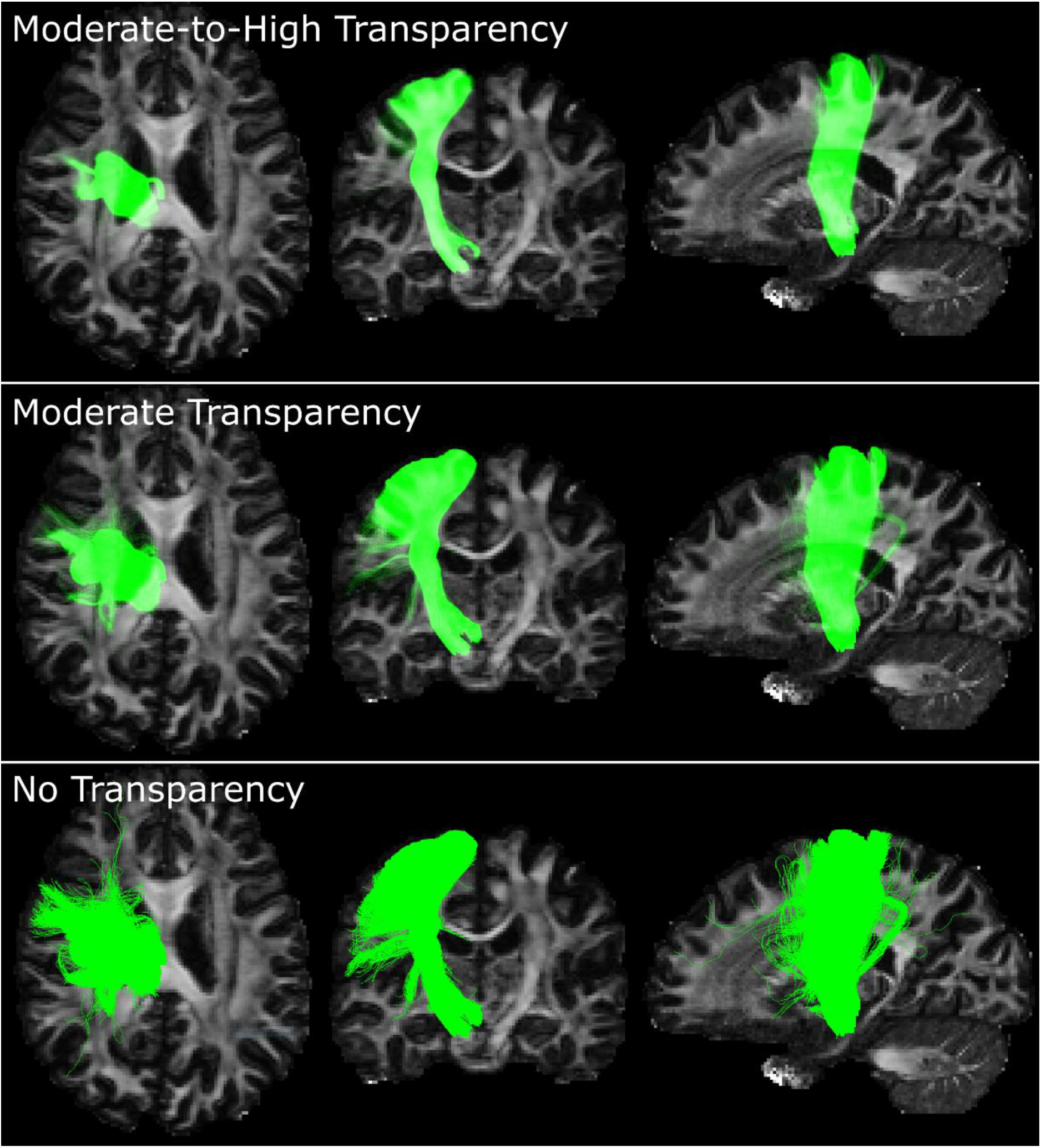
Altering transparency of streamlines can help to qualitatively judge the anatomical boundaries of a tract but requires considerable experience for use. A 100,000 streamline tractogram of the corticospinal tract is shown at moderate-to-high transparency (top row), moderate transparency (middle row) and without transparency (bottom row). Without transparency, the true boundaries are obfuscated by false-positive streamlines. With increasing transparency, the visual effect of these are reduced, but this also causes thinning of the central shaft and the disappearance of streamlines to the lateral pre-central gyrus.

In this work we explore several factors influencing tractography reproducibility and propose two methods. The first automatically estimates the number of streamlines required to achieve reliable microstructural measurements, whilst the second estimates the number of streamlines required to achieve a reproducible binarised trackmap. Both methods can be applied either prospectively, to ensure adequate streamline numbers are generated, or retrospectively, to check historical results. Theoretically, these can be applied to any desired anatomy, diffusion dataset, diffusion model, or probabilistic tractography algorithm.

## 2 Methods

We propose two metrics for determining the reliability of a tractogram. The first is a simple method that estimates the number of streamlines required to reliably sample a microstructural measure, such as FA or mean diffusivity (MD). The second is a more complex method we term *Tractogram Bootstrapping* which estimates the number of streamlines required to generate a binarised trackmap that has a known margin of error in terms of voxels included and excluded. Both methods were tested for three tracts: the corticospinal tract, the forceps major, and the long segment of the arcuate fasciculus. Implementations of both methods can be downloaded from https://bitbucket.csiro.au/projects/CONSULT/repos/tractography-reliability/. Symbols are defined in Table 1.

**Table 1.** Abbreviated terms used to describe approaches proposed in this work.

|  |  |
| --- | --- |
| AF | Long segment of the arcuate fasciculus |
| CST 1.25 | Corticospinal tract delineated at 1.25mm resolution |
| CST 2 | Corticospinal tract delineated at 2mm resolution |
| $D$ | Dice coefficient |
| $D_{0.05}(n_{samp})$ | 5<sup>th</sup> percentile of Dice coefficients for tractograms containing $n_{samp}$ streamlines |
| $D_t$ | The target Dice coefficient. If the given tract was generated twice using identical parameters, and both converted to binary trackmaps, we desire a 95% chance that the Dice coefficient between these two maps is at least $D_t$ |
| HCP | Human Connectome Project |
| FA | Fractional Anisotropy |
| FM | Forceps Major |
| MD | Mean Diffusivity |
| $n_{req}$ | The number of unique streamlines required to achieve the target reproducibility |
| $n_{tract}$ | The number of unique streamlines currently generated |
| $n_{samp}$ | The number of streamlines sampled from the unique set. This value may be greater than $n_{tract}$ if such sampling allows duplicates |
| ROI | Region of interest |
| $t_{bin}$ | Binarisation threshold |
| TB | Tractography bootstrapping |
| $W$ | Desired width of the 95% confidence interval |

### 2.1 Diffusion Metric Reliability Estimation

It is common to use a tractogram to sample from an image containing microstructural information, such as FA. One common method to achieve this is to take the value from the image at each streamline vertex (stepping coordinate), average these into a single value per streamline, and take the mean of these streamlines values to get a final average. Generating and utilising tractograms in this way is arguably a complex form of sampling, and so the more streamlines are acquired, the more reliable (though not necessarily accurate) the tractogram diffusion metric will be. As this is an average-of-averages with a large number of data points, it can be expected to generate normally distributed values, in accordance with the Central Limit Theorem (13). Thus, if a partially-complete tractogram is available, the number of streamlines required to achieve a desired margin of error can be calculated using the standard power analysis calculation (14):

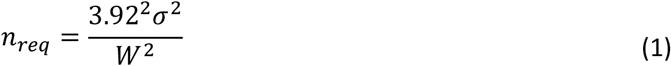

where *n_req_* is the number of required streamlines, σ is the standard deviation of values sampled from a target image using the partially generated tractogram (e.g. FA values), and *W* is the desired width of the 95% confidence interval.

We prospectively calculated the required number of streamlines to achieve a microstructural measurement of known reliability (*W*). This process is described below and summarized in Figure 3; refer to Table 1 for abbreviations. First, one thousand streamlines were generated, followed by sampling of the microstructural image. From this sample, *n_req_* was derived using Equation (1). If the number of streamlines currently generated (*n_tract_*) was greater than *n_req_*, the process exited. If not, additional streamlines were generated and appended to the tractogram. The number of additional streamlines was chosen to be *n_req_* – *n_tract_*, but constrained to the range of 1,000 to 5,000. The process then returned to the estimation of *n_req_*. The minimum (1,000) and maximum (5,000) step sizes used here were not strictly required, but solely used to improve the efficiency of streamline generation due to the large number of tractograms generated for this study. Specifically, the maximum step size reduced the risk of generating more streamlines than required. This often occurs during earlier iterations in which the algorithm overestimates *n_req_*, thus generating more streamlines than necessary. The minimum step size, by contrast, aimed to reduce the overhead of excessively stopping and starting tractography, which can occur during later iterations when small step sizes are used.

**Figure 3.**
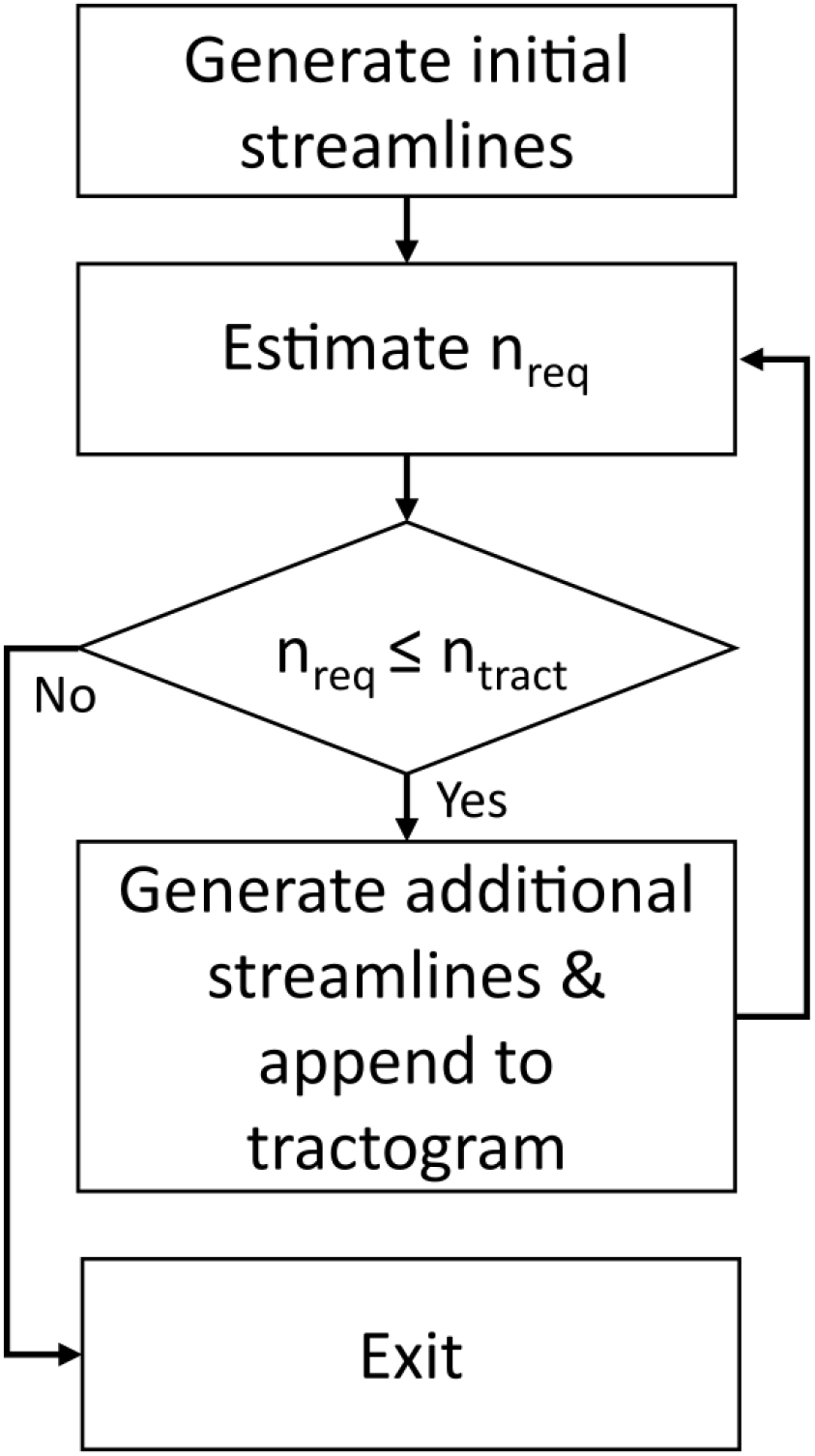
Method to estimate the required number of streamlines to achieve reliable tractography prospectively (applicable to both proposed algorithms). See text for details. Abbreviations: *n_req_*, the number of streamlines required to achieve target reproducibility; *n_tract_*, the number of streamlines currently generated.

We note that there are two alternative sampling methods. The first is to average microstructural values vertex-wise, rather than streamline-wise. This method can be used with the current procedure by providing vertex-wise (rather than streamline-wise) diffusion metrics to the proposed algorithm, and dividing the resulting *n_req_* by the mean number of vertices per streamline. The second approach is to convert a tractogram into a trackmap, binarise this, and take the average microstructural value within this region of interest (ROI). For this second approach, we refer readers to Tractogram Bootstrapping (described below), which calculates the number of streamlines required to achieve a stable binarised trackmap.

### 2.2 Tractogram Bootstrapping

Tractogram Bootstrapping (TB) was created for improving the reliability of morphological measurements, particularly with neurosurgical planning in mind. Such planning typically delineates a binary mask describing a region to be avoided for safety reasons. This safety region may be automatically generated using tractography (i.e. a binarised trackmap in the case of probabilistic tractography). As this ROI is binary, its quality can be summarized using the Dice coefficient (15), similar to traditional tissue segmentation problems. Unlike performance estimates of such traditional problems, however, the ground truth is not available, meaning that quantitative assessments of this tractography-defined region must focus on reliability rather than accuracy. Reflecting this, TB reports the number of streamlines to achieve a 95% chance that, if tracking were performed twice with identical parameters on the same data, the Dice coefficient between the two binarised trackmaps would be at least a user-defined target value (*D_t_*). For example, if *D_t_* is set to 0.9, TB would estimate the number of streamlines required such that, if tractography were performed twice, the two tractograms would have a 95% chance of a Dice coefficient of at least 0.9.

In addition to *D_t_*, TB requires two parameters that describe how binary trackmaps are generated: the trackmap voxel size, and a binarisation threshold (*t_bin_*) expressed as a fraction of the number of streamlines contributing to the map (*n_samp_*). The binarisation threshold is used to reject voxels passed through by very few streamlines. For example, a binarisation threshold of 0.001 would mean that a voxel must contain 0.001 × *n_samp_* streamlines in order to be included in the binary map.

Tractogram bootstrapping contains four major steps: sampling, similarity estimation, 5^th^ percentile calculation, and required streamline count estimation. These steps are summarized in Figure 4 and explained in detail below. Consider a tractogram with *n_tract_* streamlines. Twenty values of *n_samp_* are selected, evenly spaced from 100 to max {*n_tract_*, 10/*t_bin_*}. These lower and upper bounds were selected because initial testing suggested that failure to do so could result in overestimations of tracking reliability (see thresholding effects in Results). Sampling, similarity estimation, and 5^th^ percentile calculation steps are performed for each value of *n_samp_*, estimating reproducibility for a range of streamline counts. The final step combines these results to calculate the required streamline count (*n_req_*) for a desired level of reproducibility (*D_t_*).

**Figure 4.**
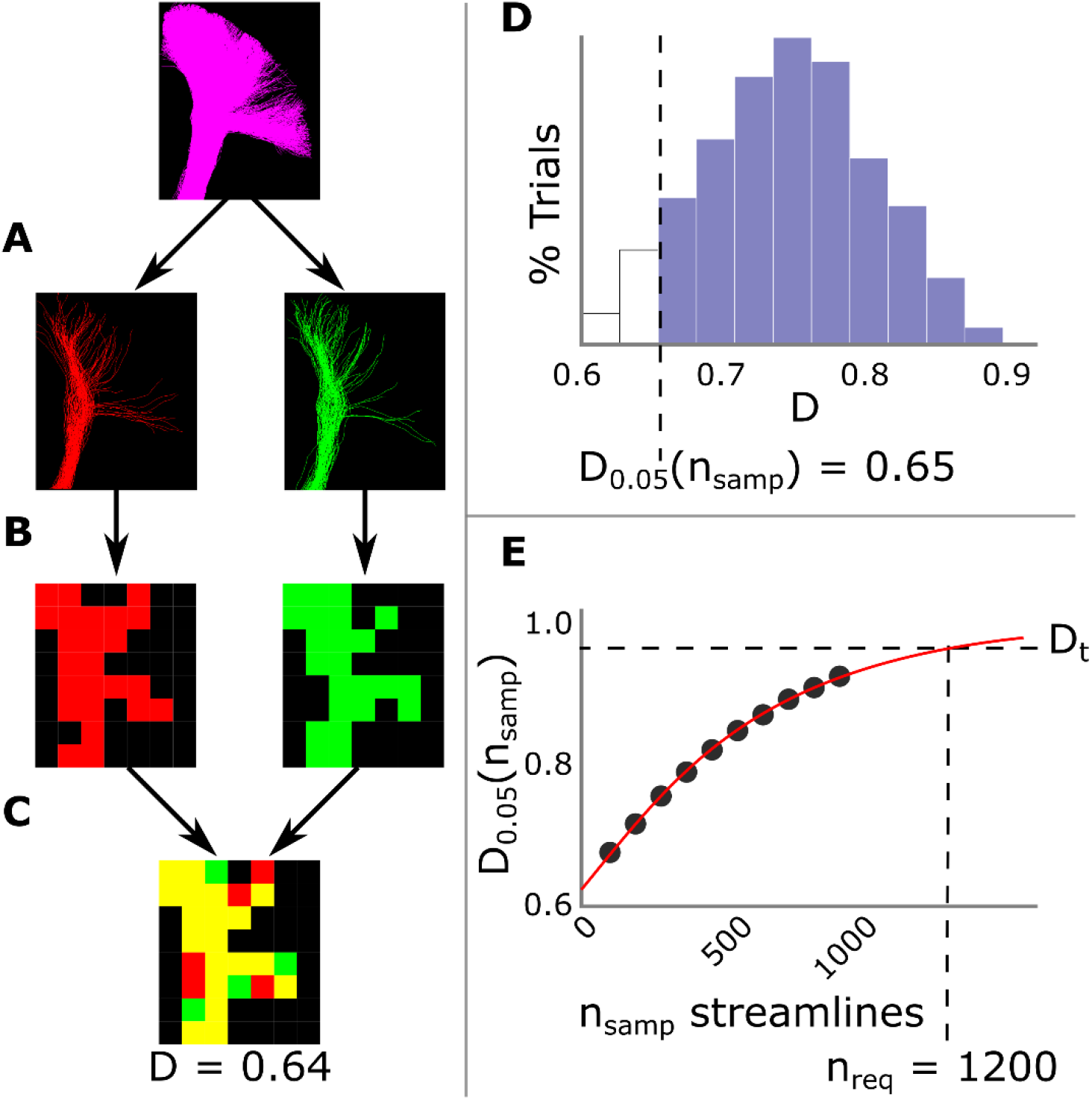
Steps performed during TB. Steps A-D are performed for each value of *n_samp_*. A – C: Sampling and Similarity Estimation (see text). D: Steps A – C are repeated 100 times, estimating Dice coefficients 100 times, shown here as a histogram. The 5^th^ percentile (*D*_0.05_(*n_samp_*)) is then calculated. E: The relationship between *n_samp_* and *D*_0.05_(*n_samp_*) is modelled via a logistic regression (red line), allowing calculation of the number of required streamlines (*n_req_*) to achieve a user specified degree of reliability (*D_t_*). Abbreviations: *D*, the Dice coefficient; *D*_0.05_(*n_samp_*), the 5^th^ percentile of Dice coefficients for tractograms containing *n_samp_* streamlines; *n_req_*, the number of streamlines required to achieve target reproducibility; *n_samp_*, the number of streamlines sampled from the tractogram.

#### 2.2.1 Sampling

A tractogram containing *n_samp_* streamlines is generated. For each value of *n_samp_*, streamlines are sampled randomly from the tractogram to generate 100 pairs of *n_samp_*-streamline tractograms (Figure 4A). Two sampling methods are used, depending on the value of *n_samp_*. When *n_samp_* ≤ 0.5 × *n_tract_*, streamlines are sampled *without* replacement (i.e. so that within a pair, no streamline appears twice). When *n_samp_* is larger, sampling with replacement (i.e. bootstrapped sampling) is performed, enabling larger samples but carrying the drawback that a given pair of tractograms may contain duplicate streamlines both within and between one another; this drawback is further described below. This combination of sampling methods was used because initial testing suggested that such an approach generally allowed *n_req_* to be estimated more accurately than sampling without replacement alone when fewer than ~2 × *n_req_* streamlines had been generated.

#### 2.2.2 Similarity Estimation

Each tractogram is converted into a trackmap that is thresholded at *t_bin_* × *n_samp_* and binarised (Figure 4B). For each pair of tractograms, the Dice coefficient of the two trackmaps (*D*) is then calculated (Figure 4C).

#### 2.2.3 Fifth Percentile Calculation

Once 100 Dice coefficients have been calculated for a particular *n_samp_*, the 5th percentile of this metric, denoted here as *D*_0.05_(*n_samp_*), is calculated (Figure 4D). *D*_0.05_(*n_samp_*) is an estimation of expected reproducibility of future tractography generation for a particular streamline count. Specifically, if we were to generate two new tractograms, each containing *n_samp_* streamlines, we have an approximately 95% chance that the Dice coefficient between these would be at least *D*_0.05_(*n_samp_*).

Unlike sampling without replacement, bootstrapping is prone to inflation of Dice coefficients, and thus *D*_0.05_ estimates. Such bias can be calculated by performing both bootstrapping and sampling-without-replacement where possible. This knowledge can be used to correct bias where only bootstrapping is possible. Due to metric discontinuities induced by thresholding (see Results), however, this is only possible for very few values of *n_samp_*, and sometimes not at all. In the interest of brevity, we refer interested readers to Supplementary Materials for an explanation of how this correction was performed.

#### 2.2.4 Required Streamline Count Estimation

The final step (Figure 4E) is performed once all *D*_0.05_(*n_samp_*) values have been calculated. To predict the number of streamlines required to meet criteria *D_t_* (i.e. the minimum *n_samp_* such that *D*_0.05_(*n_samp_*) ≥ *D_t_*), the relationship between *D*_0.05_ and *n_samp_* can be described by a traditional four-parameter logistic curve whose inflection point is fixed at one streamline and remaining parameters *a*, *b* and *d* are estimated using the Levenberg-Marquardt technique:

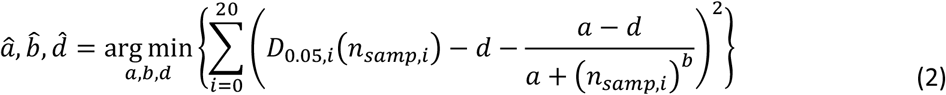

The number of streamlines required to achieve the user-specified confidence criteria *D_t_* can then be estimated from the inequality below for the streamline number:

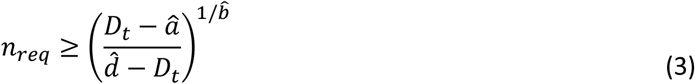

Note that during development simpler alternatives to Equation 2 were explored but demonstrated substantially poorer fits to data.

### 2.3 Comparison with Cross Validation

Cross validation was used to assess the ability of the two proposed algorithms to estimate *n_req_* for a range of reliability criteria and white matter tracts. We utilized the first 40 ‘minimally pre-processed’ diffusion datasets from the Human Connectome Project (HCP) Young Adult dataset (1200 Subjects Release) (16). For each dataset, during tracking we prospectively estimated the number of streamlines required to meet the criteria in question, using the previously described methods and the process shown in Figure 3. For each dataset and criterion, we then generated an additional 100 tractograms each containing the predicted number of required streamlines. These 100 additional tractograms were compared to one another (in terms of diffusion metrics or similarity) to ascertain the *actual* reliability of this tractography.

#### 2.3.1 Image Processing

Diffusion images were used in their ‘minimally preprocessed’ state, as provided in the HCP 1200 Subjects Data Release (unique directions: 90 @ 1000 s/mm^2^, 90 @ 2000 s/mm^2^, and 90 @ 3000 s/mm^2^, plus 18 @ b=0 s/mm^2^). This minimal preprocessing included correction for b0 intensity inhomogeneities, EPI distortion, eddy currents, head motion, gradient non-linearities, as well as reorientation and resampling to 1.25mm isotropic (17). Each diffusion scan contributed to three datasets: a high-resolution multishell dataset containing unaltered images; a ‘downsampled multishell’ dataset generated by downsampling preprocessed images to 2mm isotropic; and a single-shell dataset generated by removing all but 50 volumes from the downsampled multishell dataset (5 @ b=0 s/mm^2^; 45 @ b=1000 s/mm^2^, selected to be approximately evenly distributed on the sphere using code provided in the aforementioned git repository). The single-shell dataset consisted of the b=1000 s/mm^2^ shell so that the tensor images would be maximally similar between the three datasets, as these were calculated from this shell in all instances. Fiber orientation distribution images were generated using MRtrix3’s (18) multi-shell multi-tissue constrained spherical deconvolution method (multishell data) or Single-Shell 3-Tissue constrained spherical deconvolution (single-shell dataset; https://3Tissue.github.io), in conjunction with the Dhollander algorithm to estimate the tissue response functions (19,20).

We generated tractograms of the right corticospinal tract for all three datasets to observe the effects of spatial and angular resolution. To also observe the effects of anatomy, we also generated tractograms for the long segment of the right arcuate fasciculus and the forceps major using the multishell downsampled dataset. The multishell downsampled dataset was chosen for this task to reduce computational overhead and to test the proposed algorithms at a resolution more typically seen in current literature.

High resolution (0.7 mm isotropic) structural T1 MPRAGE images were denoised using Global Approximate Block Matching (21). The registration between the T1 and diffusion data set was ensured by performing a rigid registration between the T1 and first b=0 image of the series, using ANTS. ANTS SyN (22) was then used to calculate the non-rigid registration between the result and the MNI ICBM 152 template (23) to enable the later transfer of ROIs from MNI space into diffusion space.

#### 2.3.2 Tractography

For tractography we used MRtrix3’s iFOD2 algorithm (24). Unless specified, ROIs were those defined by the Freesurfer-based parcellation provided in the HCP dataset. Other described ROIs are supplied as figures in Supplementary Materials.

The right corticospinal tract was seeded from the grey matter / white matter boundary of the right precentral gyrus to the brainstem mask. The corpus callosum mask was dilated by one voxel (18-connected) and used as an exclusion mask.

The long segment of the arcuate fasciculus was seeded from grey matter / white matter boundary found within the pars opercularis. The grey matter / white matter boundary of the superior temporal lobe posterior to MNI *y* = 14.5*mm* acted as an inclusion mask. Manually delineated exclusion and inclusion masks (Supplementary Figures 4 – 6), designed to reduce anatomically implausible streamlines, were moved from MNI space into diffusion space. The aforementioned dilated corpus callosum mask formed a second exclusion mask.

The forceps major was tracked both from left-to-right (50% of streamlines), and from right-to-left, the results of which were combined to form a final tractogram. Seed or inclusion masks in each hemisphere consisted of the lateral occipital lobe, cuneus and pericalcarine fissure. The splenium was an additional inclusion mask in both cases. An exclusion mask (Supplementary Figure 7), manually delineated on the MNI template and moved into each subject’s diffusion space, reduced anatomically implausible streamlines.

#### 2.3.3 Cross Validation

To test the tractography metric reliability method, the following was performed for each type of tract, in each subject, targeting standard deviations of 0.001 for FA measurements and 10^−6^ for MD measurements. Initially, a tractogram was generated using the methodology summarized in Figure 3. To ensure a fair assessment of this algorithm, if this method generated more than *n_req_* streamlines (i.e. overshot due to the minimum number generated on each iteration), streamlines were removed such that the streamline count was *n_req_*. A further 100 tractograms were then generated in the normal manner, each with *n_req_* streamlines. The mean FA or MD measurement was taken from each of these 100 tractograms using MRtrix3 and the standard deviation for each subject was compared with the specified stopping criteria.

Tractogram bootstrapping was tested for the same anatomical tracts for a range of confidence parameters, listed in Table 2. We note that Conditions A1 and A2 are too lenient for neurosurgical applications and were only used here to explore the robustness of the proposed algorithm. A tractogram was generated using the process described earlier, until the stopping criterion was met, and the streamline count restricted to *n_req_* in the case of an overshoot. One hundred additional tractograms with *n_req_* streamline counts were then generated, converted into binary trackmaps, and paired into 50 sets of two. For each pair, the Dice coefficient was calculated in the way previously described. The 5^th^ percentiles of these proportions were then recorded and compared with the appropriate *D_t_* value.

**Table 2.** Parameters for the conditions tested. Track-map resolution matched the diffusion image resolution (1.25mm or 2mm isotropic). Abbreviations: *D_t_*, target dice coefficient; *t_bin_*, binarisation threshold.

| Condition | $D_t$ | $t_{bin}$ |
| --- | --- | --- |
| A1 | 0.9 | 0.01 |
| B1 | 0.95 | 0.01 |
| C1 | 0.97 | 0.01 |
| A2 | 0.9 | 0.001 |
| B2 | 0.95 | 0.001 |
| C2 | 0.97 | 0.001 |

## 3 Results

### 3.1 Tractography Metric Reliability Estimation

The number of streamlines required for reliable microstructural measurements varied considerably by type of microstructural measurement (i.e. FA or MD), anatomical tract, and dataset (Figure 5, top panel). Of particular note, the number of streamlines required to achieve reliable MD measurements varied by over two orders of magnitude, depending on the anatomical tract and dataset in question (arcuate fasciculus minimum, 107; forceps major maximum, 30100). At these *n_req_* values, cross validation demonstrated actual standard deviations similar to target values for FA and MD in all anatomical tracts targeted (Figure 5, middle panel). These standard deviations did not differ between the five tracking conditions (one-way ANOVAs; both with *p* > 0.4). When we purposefully selected fewer than *n_req_* streamlines, standard deviations were larger than target values (Figure 5, bottom panel).

**Figure 5.**
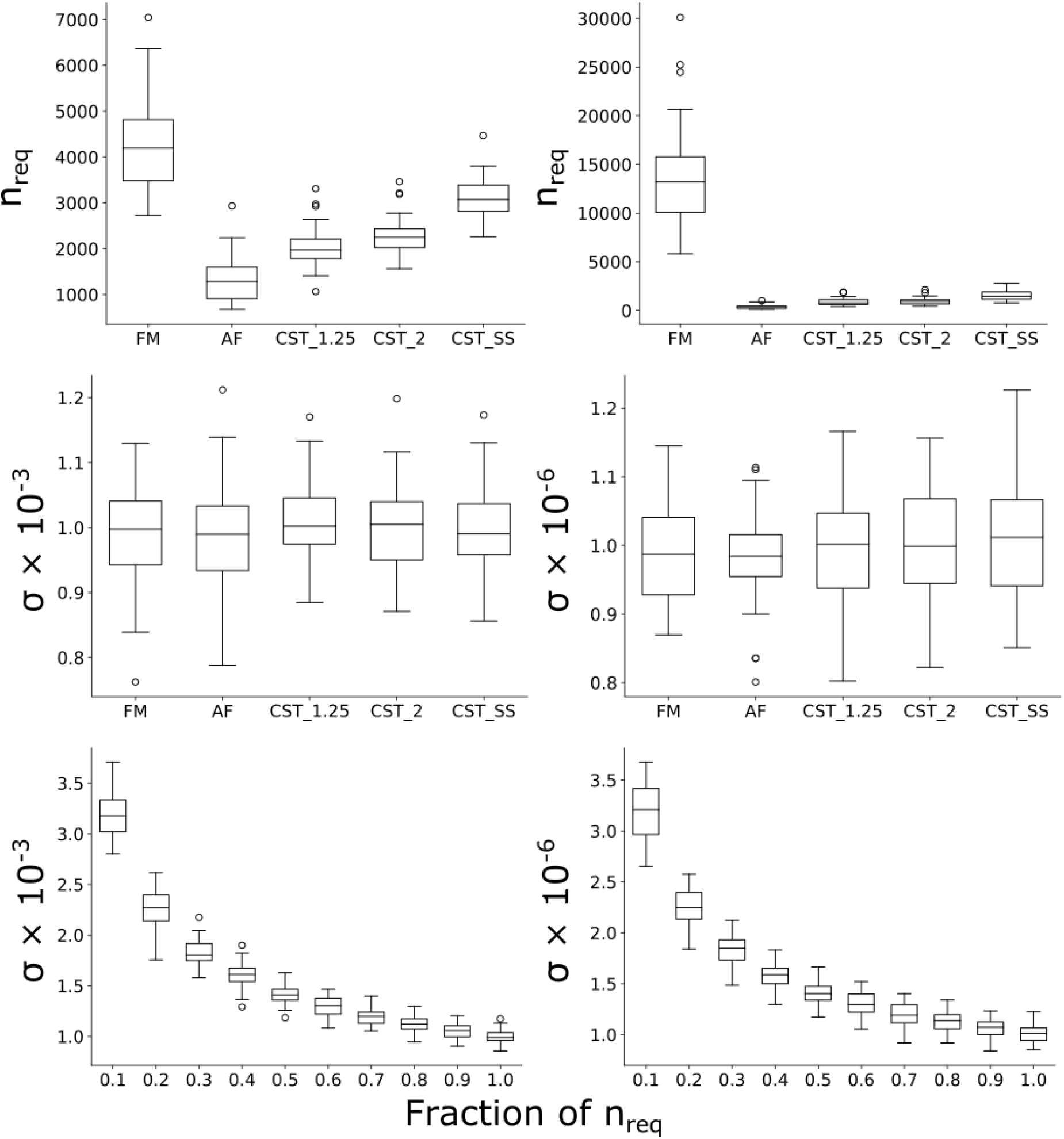
Number of streamlines required and errors in achieving target standard deviations of 10^−3^ (FA, left column) and 10^−6^ (MD, right column). Each participant contributed a single datapoint to each box in each plot. Top: Predicted number of streamlines required to achieve target reliabilities. The number of required streamlines varied strongly depending on the anatomy and microstructural measure in question. Middle: The actual standard deviations at *n_req_*, as calculated by cross-validation. In all instances, actual standard deviations were close to that of the target values. Bottom: The actual standard deviations at fractions of *n_req_*, as calculated by cross-validation, pooled across all participants and anatomical tracts. Abbreviations: AF, arcuate fasciculus; CST 1.25, corticospinal tract at 1.25mm resolution; CST 2, corticospinal tract at 2mm resolution; CST SS corticospinal tract at 2mm resolution with single shell data; FM, forceps major; *n_req_*, the number of streamlines required to achieve target reproducibility.

### 3.2 Tractogram Bootstrapping

#### 3.2.1 General Observations

Regardless as to the anatomy in question, when binarisation was performed without thresholding, the volumes of the resulting trackmaps increased in a logarithmic manner (Figure 6). When thresholding was applied as a function of streamline count, these volumes reached a plateau if sufficient streamlines were generated. However, this trend demonstrated discontinuities when this threshold reached the next integer (e.g. at 1000, 2000, 3000 streamlines), the influence of such discontinuities diminishing as streamline count increased. In some instances, this meant that when binarised trackmaps where generated from fewer streamlines, their volumes would be higher than when generated with much larger numbers of streamlines (Figure 6, Figure 7). We also experimented with an alternative strategy (4), where the threshold was set at 1% of the maximum trackmap intensity. This method showed the same behavior, with the additional drawback that the streamline counts at which these discontinuities would occur was not easily predictable, varying by dataset and tract type.

**Figure 6.**
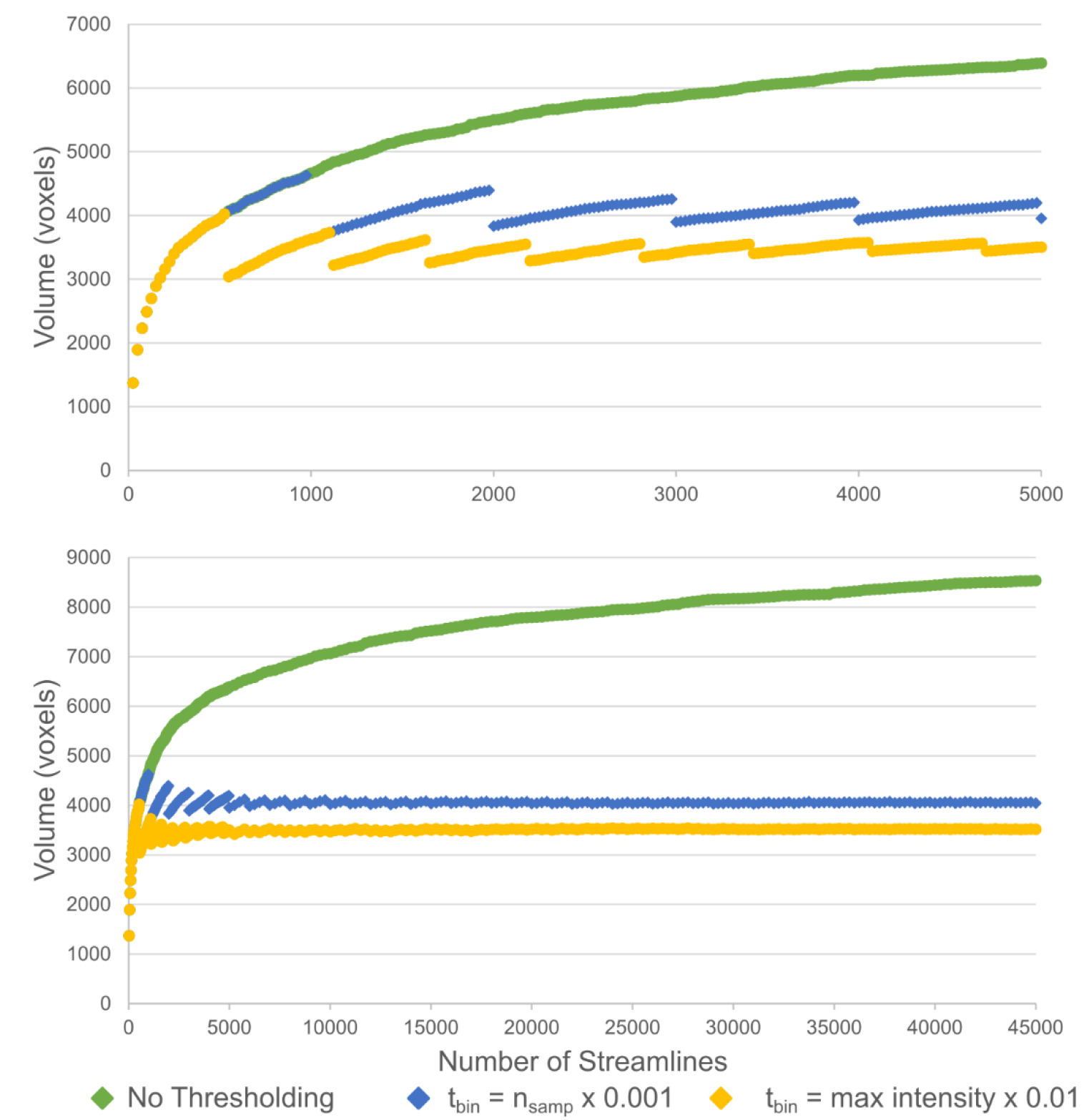
Relationship between binarised trackmap volume and number of streamlines. Upper and lower graphs are the same data shown at two different x-axis ranges. These data were generated by tracking the forceps major of an HCP participant included in the current study. The three lines represent the binarised trackmap volume when not thresholding (green, top), thresholding at 0.001 x streamline count (blue, central), and at 0.01 x the maximum trackmap intensity (gold, bottom). Abbreviations: *n_samp_*, the number of streamlines sampled from the tractogram; *t_bin_*, the binarisation threshold.

**Figure 7.**
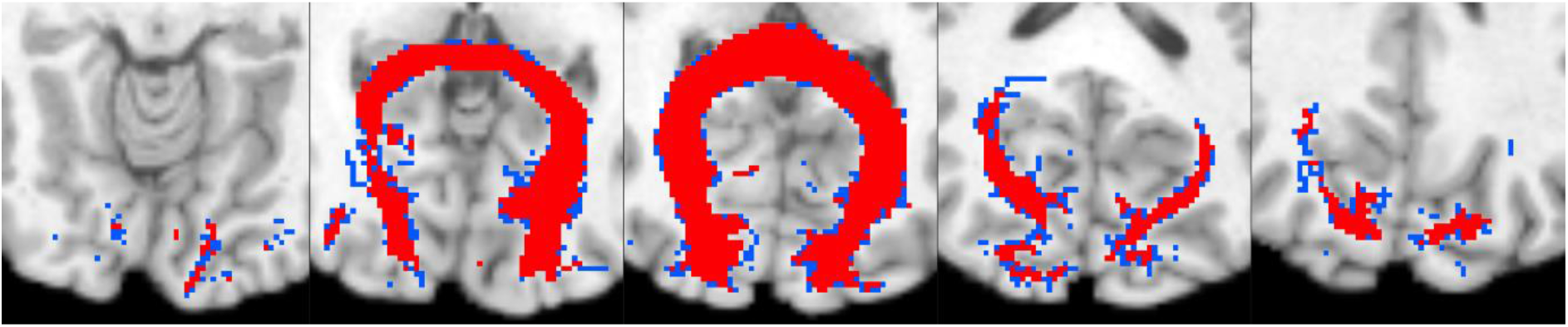
Five axial slices of a forceps major trackmap, thresholded at 0.001 x the streamline count and binarised. Blue and red together indicate the binary map when 999 streamlines are available. Red alone shows the binary map when an additional two streamlines were added to this tractogram and thresholding was performed using the same rule. Notice the high number of voxels removed (blue) due to this minute increase in streamline number raising the threshold to the next integer value.

#### 3.2.2 Predictive Performance

The predicted number of streamlines differed substantially depending on the dataset, anatomy to delineate, and target reproducibility (Figure 8, Left). To meet the reproducibility criteria at a resolution of 2mm with multishell data, the number of required streamlines (*n_req_*) ranged from 593 (Condition A2; arcuate fasciculus) to 16738 (Condition C2; forceps major). The number of streamlines required to delineate the corticospinal tract was also substantially higher at a resolution of 1.25mm than at 2mm, but higher still for the 2mm resolution single-shell dataset.

**Figure 8.**
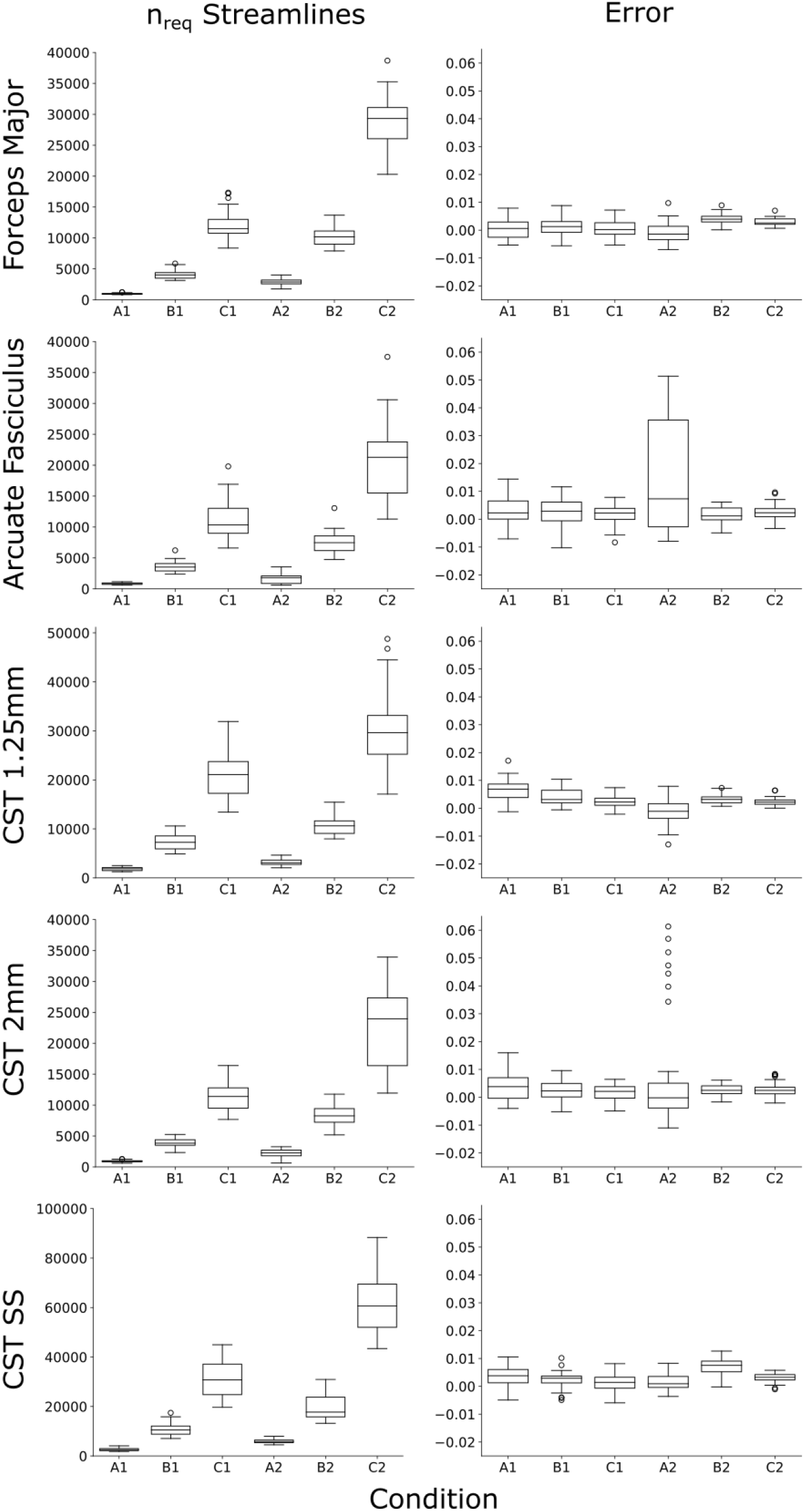
Left: The required number of streamlines (*n_req_*), as calculated by tractogram bootstrapping for different anatomy and reproducibility criteria. Note the CST conditions have different y-axis ranges. Note how *n_req_* differs clearly by the anatomy, reproducibility condition, spatial resolution and angular resolution. Right: the amount of error, expressed as *D_t_* – *D_actual_*, where these actual values were assessed by cross validation using *n_req_* streamlines. In most circumstances, actual error was within 1% of that desired. Both arcuate fasciculus and forceps major delineations were at a 2mm isotropic resolution. Each participant contributed a single datapoint to each box in each plot. Abbreviations: CST 1.25, corticospinal tract at 1.25mm resolution; CST 2, corticospinal tract at 2mm resolution; CST SS corticospinal tract at 2mm resolution with single shell data; A1, A2, B1, B2, C1, C2: conditions tested, as defined in Table 2; *D_t_* the target Dice coefficient; *D_actual_*, the actual dice coefficient.

The actual Dice coefficient at *n_req_*, as assessed by cross validation, differed by less than 0.01 from target values (*D_t_*) across conditions B1, B2, C1, and C2 in 99% of all tests (Figure 8, Right). Such absolute error was below 0.01 in 90% of cases for conditions A1 and A2. When purposefully selecting fewer than *n_req_* streamlines, Dice coefficients were lower than *D_t_* for all tracts and conditions (Figure 9, Supplementary Materials S3).

**Figure 9.**
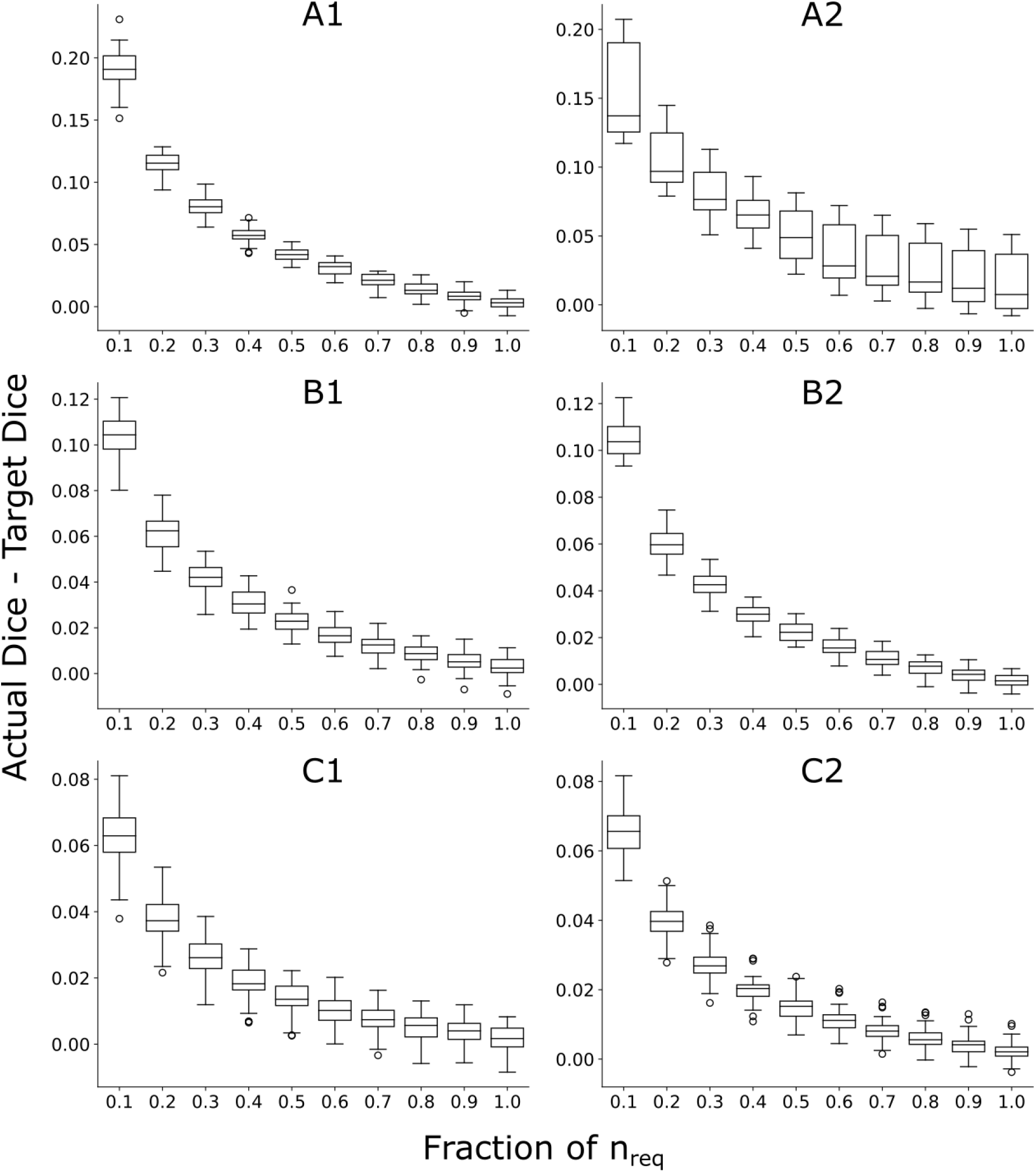
Relationship between streamline number, normalized to *n_req_*, and tractography reliability of the arcuate fasciculus. The y-axis indicates the actual Dice coefficient (*D_actual_*) minus the target Dice coefficient (*D_t_*). Each participant contributed one datapoint to each box, in each plot. A1, A2, B1, B2, C1, and C2 refer to different stopping criteria, defined in Table 2. Error progressively reduced as more streamlines were added, reaching approximately zero for most stopping criteria when *n_req_* streamlines had been generated. Relationships for other anatomical tracts were similar and can be found in Supplementary Materials.

## 4 Discussion

Tractography is utilized in both clinical and scientific contexts for the purposes of taking morphological and microstructural measurements. In both contexts, generating a sufficiently high number of streamlines is critical to ensuring that such measurements are reproducible. Tractography can be a computationally expensive process in terms of generation, viewing, and storage. Choosing a practical number of streamlines is not a simple task, because the relationship between streamline count and reproducibility is likely to depend on a great number of patient and image-related factors. Here, we proposed two methods designed to automatically calculate the number of streamlines needed for reliable tractography in an individual dataset. Both methods can be performed prospectively. A major benefit to this approach is that it can both prevent inflation of Type I and II errors due to insufficient streamline generation, as well as avoid excessive streamline generation that can be computationally expensive.

We demonstrated how standard statistics can be utilized to estimate how many streamlines are required to achieve reliable microstructural measurements, such as FA or MD. When used prospectively, this approach reliably generated tractograms that gave FA or MD measurements with true margins of error close to the targeted margin of error (Figure 5). Vastly different numbers of streamlines were required for different anatomical tracts and microstructural measures (Figure 5). Presumably, such differences were due to different tract types having different delineation reliabilities, as seen in our experiments on trackmap reliability, as well as different distributions of microstructural values throughout their volume (e.g. differing proportions of voxels containing crossing fibers or partial volume effects with ventricular CSF). In the past, it has been common to base streamline counts on qualitative assessments (such as the appearance of a test tract) or default software values, rather than by considering the microstructural measures which are intended to be sampled. In the present study, the large difference between required numbers of streamlines for FA and MD in the forceps major highlights that such an approach is unlikely to fairly assess how many streamlines are required for reliable measurements. The number of required streamlines in several test cases here also demonstrated that streamline counts in the low thousands, sometimes considered to be sensible or even excessive, might be inappropriate for some datasets and hypotheses. Given the variability demonstrated, we wish to make clear that it is not appropriate to utilize the estimates reported here to choose streamlines counts in other datasets. Rather, we encourage readers to apply the methods provided here to their own datasets to ensure that adequate reliability is obtained.

Following this analysis, we turned our focus to the generation of binary trackmaps – ROIs generated from probabilistic tractography that can be more suited to some neurosurgical settings. An important finding is that the method by which a tractogram is binarised can markedly affect the volume of the resulting map. Specifically, if thresholding is not performed before binarisation then tract volume grows until virtually the entire brain is filled (Figure 6). The implications of this are that, when tractography is used to estimate the safety of a surgical procedure, failure to apply a threshold can result in an unreasonably large estimate of risk, potentially resulting in surgical intervention being wrongly altered or rejected over safety concerns. By contrast, voxelwise thresholding allows tract volumes to reach a plateau. However, thresholding causes discontinuities in volume at multiples of the inverse of the binarisation threshold (1/*t_bin_;* Figure 6), which appear to be particularly strong for the first and second multiples of this threshold, but decrease in amplitude with increasing numbers of streamlines. To avoid this issue, based solely on the data seen here, we caution against selecting a streamline count below four times the inverse of the binarisation threshold used. We also experimented with an alternative binarisation approach based on the maximum trackmap intensity, but this demonstrated the same problem and had an additional drawback in that predicting where discontinuities would occur would be difficult or impossible before tracking takes place. We do note that non-integer trackmaps are possible in MRtrix3 (25), which might avoid the existence of discontinuities, but caution their use on the basis that the interpretation of these trackmap values is not straightforward. Specifically, these ‘precise’ trackmaps allow each streamline to contribute values greater than one to each voxel when it passes through a voxel non-perpendicularly. This means that the resulting map no longer reflects streamline count passing through a region in a straightforward way: for example, higher values can be expected in areas of curvature or where streamlines travel at an angle relative to the voxel orientation. This makes the choice of a threshold less intuitive than the simpler mapping used here.

To achieve a reliable map, the number of required streamlines estimated by TB differed substantially depending on the participant, anatomy to delineate, binarisation threshold, spatial resolution, angular resolution, and target reproducibility (Figure 8). For realistic parameters (B1, B2, C1, and C2), these estimations appear to have been accurate: when cross validation was performed for *n_req_* streamlines, 99% of cases resulted in actual *D*_0.05_ values within 0.01 of *D_t_*. For criteria A1 and A2, this success rate was somewhat lower, potentially because the number of streamlines required was often below 100, which is the bottom limit at which the algorithm explicitly estimates *D_t_*. We emphasise again that criteria as lax as A1 and A2 should not be used, but were merely tested here to evaluate performance of tractogram bootstrapping under a range of input parameters.

We note that the present method is purposefully designed for a limited scope of applications; in other situations it may be appropriate to extend this work or to use more appropriate previously published methods. For example, the present work is targeted towards identifying, or measuring metrics from singular tracts. For whole-brain based analyses, a more sophisticated tool such as SIFT2 (26) is likely to be more appropriate to ensure streamline counts are comparable across the brain’s physical network. However, SIFT2 is not appropriate for single-tract analyses, as it relies on contextual information supplied by other tracts and cannot currently estimate the number of streamlines required to achieve reliable microstructural measurements. One potential extension of our method is to estimate *n_req_* for non-binarised trackmaps. To achieve this, it is a relatively trivial exercise to avoid binarisation and replace *D_t_* in the current implementation with an image similarity metric, such as a normalised sum of absolute differences. Although beyond the intended scope of the current work, our initial informal testing with this appears to show relatively robust results. Such metrics, however, are neither particularly interpretable nor intuitive, meaning that choosing appropriate stop criteria is potentially no less arbitrary than selecting a streamline count directly. We note that coefficient of variation is an intuitive metric that has been previously used to compare trackmaps (6) but, in our experience, can behave erratically when ‘stray’ streamlines are generated.

Finally, we reiterate that the proposed methods are solely designed to reduce variability caused by insufficient streamline counts. That is, the proposed methods do not guarantee that such tractography is accurate, simply that the streamline generation command itself provides reproducible outputs when applied to the same scan repeatedly. An interesting extension to the present work would be assessing to what extent additional factors affect the scan-rescan reproducibility of tractography and associated microstructural measurements. Some answers and solutions to issues, however, may be complex as such reproducibility is likely to depend on a wide range of currently non-standardised and interacting factors including the MR sequence, preprocessing steps, anatomy investigated, type and presence of pathology, ROI placement method, streamline generation algorithm and even gradient non-linearities of the scanner in question (27).

In conclusion, we have presented two methods. The first automatically estimates how many streamlines are required to achieve reliable microstructural measurements, whilst the second estimates how many streamlines are required to achieve a reliable binarised trackmap. When we repeatedly generated tractograms, each containing the estimated number of streamlines, we found microstructural measurements and resultant trackmaps had levels of reproducibility closely aligned to that targeted. We hope that by making these tools available (https://bitbucket.csiro.au/projects/CONSULT/repos/tractography-reliability/), researchers can more easily select the appropriate number of streamlines for their application, removing the need to rely on rules of thumb or the qualitative appearance of resultant tractograms.

## Acknowledgements

The authors are grateful to Ashley Gillman for his help in preparation of this manuscript.

