## Supplemental Materials for "How Many Streamlines are Required for Reliable Probabilistic Tractography? Solutions for Microstructural Measurements and Neurosurgical Planning"

### Supplementary Materials

#### S1. Correction of Bootstrapping

For the following text, $n_{tract}$ refers to the number of unique streamlines generated through normal means (such as MRtrix's tckgen), and $n_{samp}$ refers to the number of streamlines we wish to sample from that set to form a 'new' tractogram.

Bootstrapping is used where generation two $n_{samp}$-size tractograms is not possible by sampling without replacement (i.e. where $2n_{samp} > n_{tract}$). Bootstrapping samples *with* replacement, and so is prone to generating datasets (tractograms) containing duplicates. This means that a given resampled tractogram is likely to be more similar to itself, and to the other tractogram to which it is paired, than would be the case if it were generated from scratch using a tool such as tckgen. In the context of the method presented here, this can mean that bootstrapping results in an inflated Dice coefficient estimation, particularly when $n_{tract}$ (the number of available streamlines to sample from) is not substantially larger than $n_{samp}$ (the number of streamlines in a sample). By contrast, no such issue is likely when sampling without replacement, as tractograms to be compared lack duplicates within and between themselves.

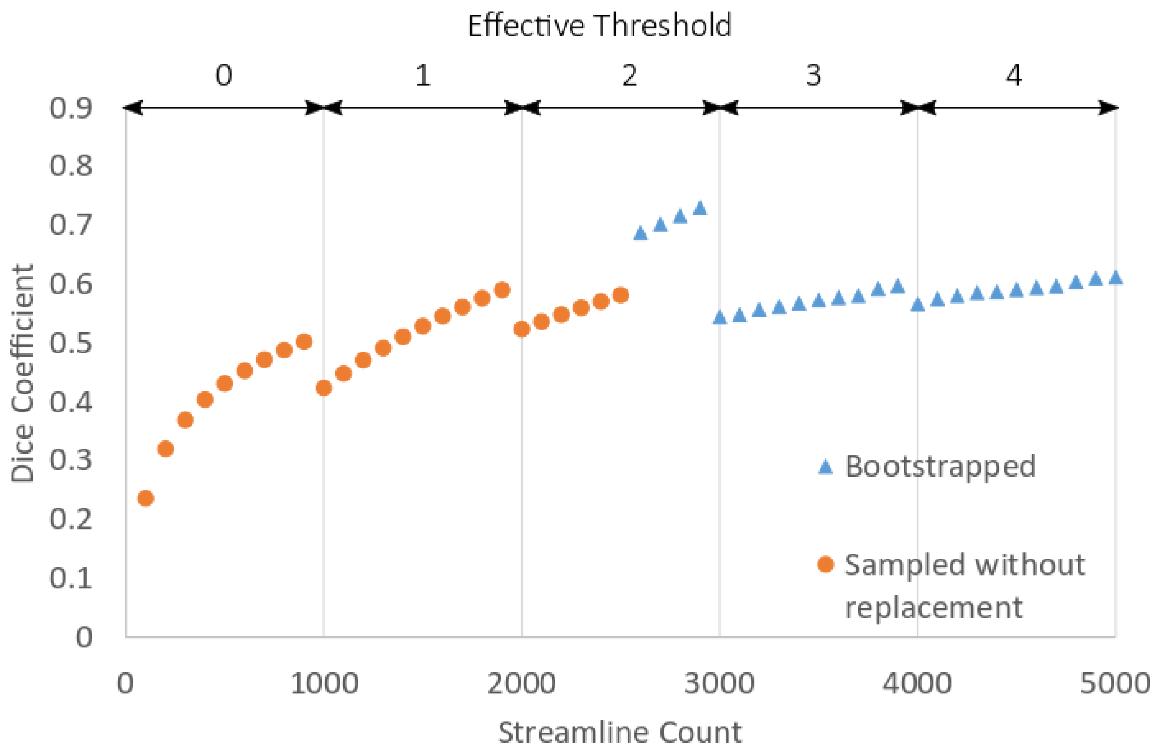


**Supplementary Figure 1. Type of sampling with example Dice Coefficients for $n_{tract}$ of 5000 and $t_{bin}$ of 0.001. Data shown is for illustration only.**

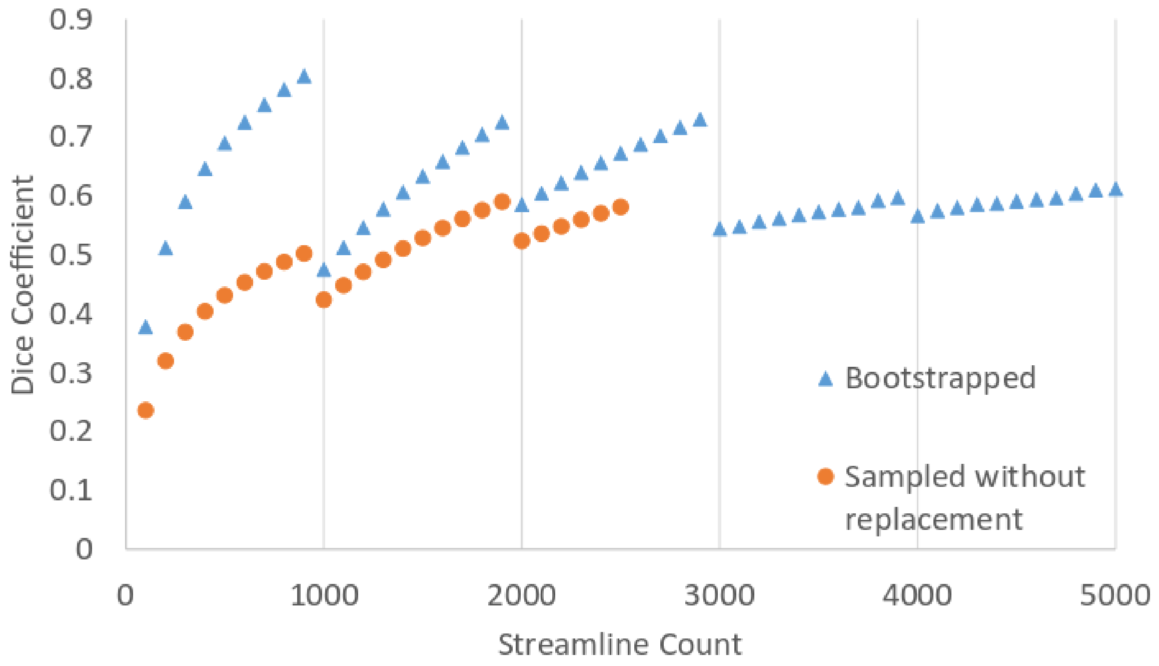


**Supplementary Figure 2. Bootstrapping can overestimate dice coefficients, compared to sampling without replacement, particularly for lower numbers of streamlines. Data shown is for illustration only.**

If bootstrapped sampling and sampling without replacement are performed for the same value of $n_{samp}$, the difference between these estimations represents bootstrapping's optimistic bias. If this is performed for a range of values of $n_{samp}$, the relationship between $n_{samp}$ and bias can then be used to predict, and to correct, the level of bias where only bootstrapping is possible (i.e. where $2n_{samp} > n_{tract}$).

The limitation of this method is that the relationship between $n_{samp}$ and Dice coefficients are discontinuous due to the effects of thresholding. This is so because trackmaps reflect streamline counts in each voxel (i.e. they contain only integers), meaning that binarisation thresholds effectively change in a stepwise manner (effectively increasing by 1 every $1/t_{bin}$ streamlines; Supplementary Figure 1). For example, for a $t_{bin}$ of 0.001, the threshold between $1 \leq n_{samp} \leq 999$ is effectively 0, and $1000 \leq n_{samp} \leq 1999$ is effectively 1. As such, for $t_{bin}$ of 0.001, the relationship between $n_{samp}$ and Dice coefficient (or bias in this measure) is smooth except for marked discontinuities every 1000-streamline increase in $n_{samp}$ (see Results). This means that estimations of bias for $1 \leq n_{samp} \leq 999$, for example, are thus not suitable for performing bias correction for values of $n_{samp} > 999$.

The consequence of these discontinuities is that correcting bias is only possible for a few values of $n_{samp}$. Concretely, for values of $n_{samp}$ where sampling without replacement is possible, bias could be estimated but bootstrapping is not actually required. For values of $n_{samp}$ with a *larger effective threshold* than all estimates performed by sampling with replacement, bias cannot be estimated and so not be corrected. However, bootstrapping bias can be corrected for remaining values of $n_{samp}$ that have the *same effective threshold* as at least two estimations performed by sampling with replacement. An example is shown in Supplementary Figures 1 - 3. In this example, $n_{tract}$ = 5000 streamlines and $t_{bin}$ = 0.001, meaning discontinuities can be expected at $n_{samp}$ = 1000, 2000, 3000, 4000, and 5000 (Supplementary Figure 1). Bootstrapped estimates are not required for $n_{samp} < 2500$, as sampling-without-replacement is possible. For $n_{samp} > 2500$, bias cannot not be ascertained as sampling without replacement is impossible. For $n_{samp}$ = 2000…2500, however, both sampling methods are possible, allowing bias to be estimated (Supplementary Figure 3, top) and bias correction to be applied to bootstrapped estimates from 2501 to 2999 inclusive. (Supplementary Figure 3, bottom)

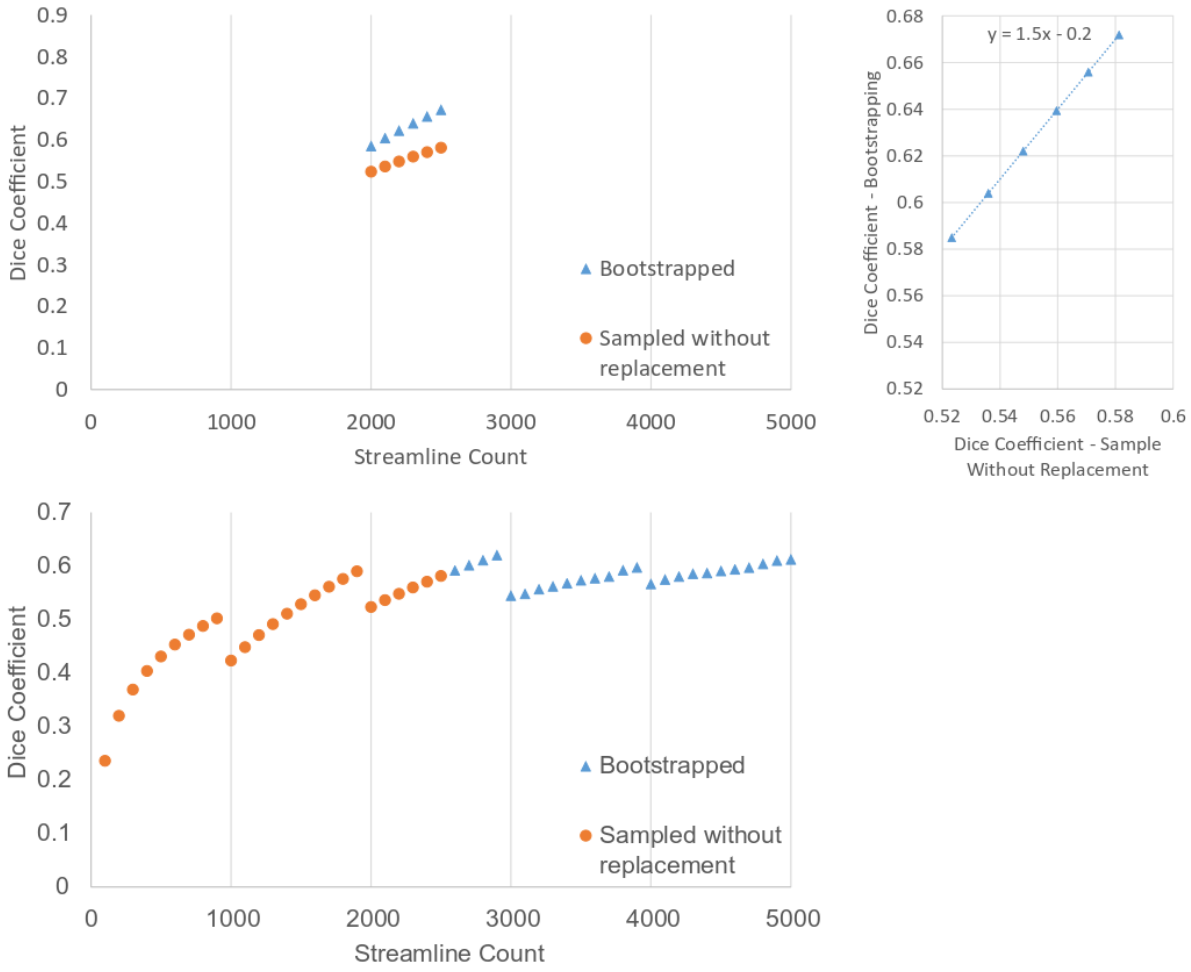


**Supplementary Figure 3. For $n_{tract}$ = 5000 and $t_{bin}$ = 0.001, bootstrapping bias can be calculated using streamline counts of 2000 – 2500 for both sampling methods (top left), by finding the linear relationship between these variables (top right). This can then be used to adjust Dice Coefficients for bootstrapped samples from 2500 < $n_{samp}$ < 3000 (bottom).**

It is noteworthy that more complex methods exist to correct bootstrapping that are less restricted to values of $n_{samp}$. In initial testing, however, methods such as the .632 bootstrap (1) markedly overcompensated bias in this particular application, potentially due to the non-smooth nature of such bias caused by binarisation.

### S2. Tractography Masks

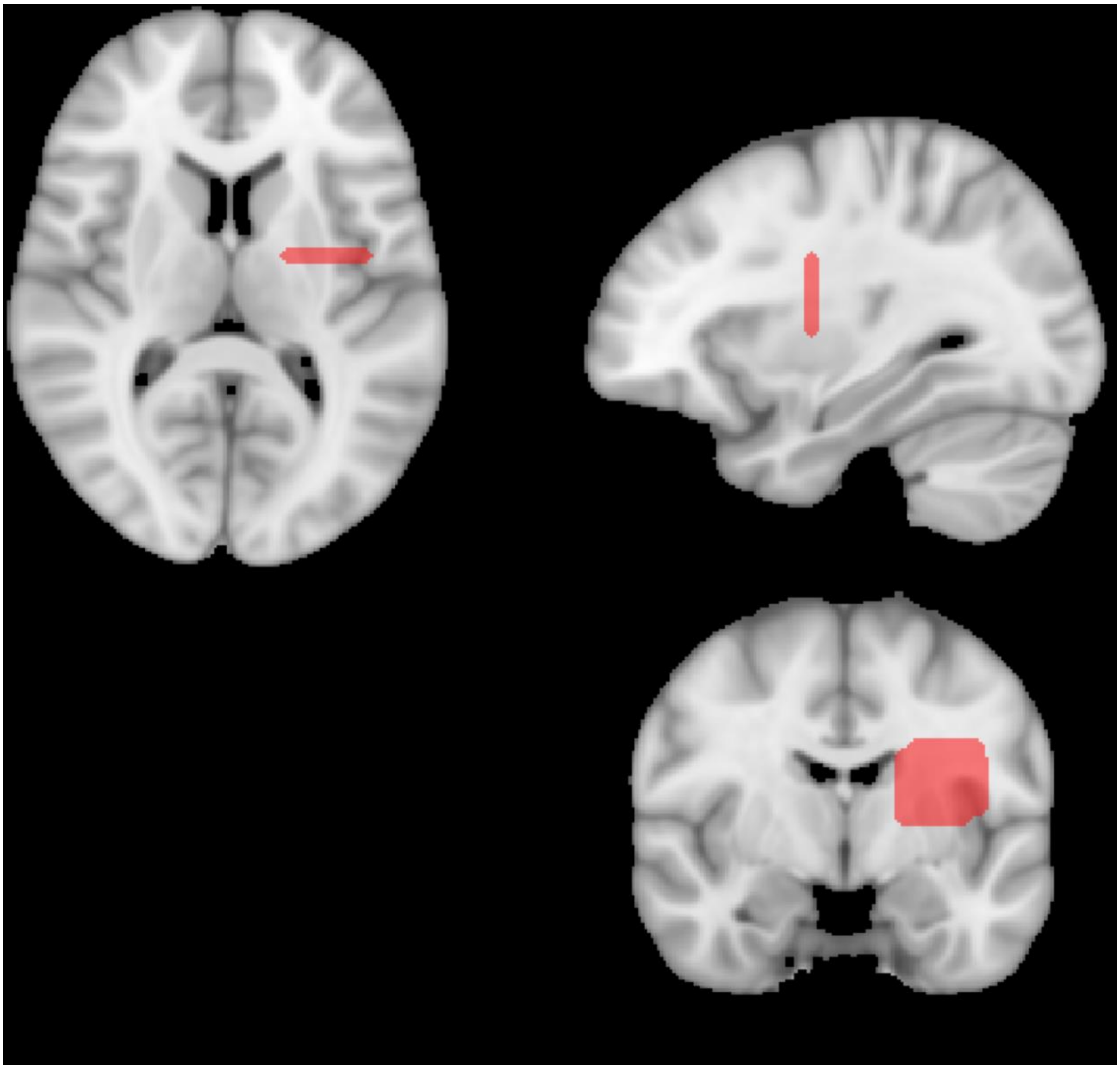

**Supplementary Figure 4. Inclusion mask for tractography of the arcuate fasciculus. The temporal lobe inclusion mask was automatically generated on a per-dataset basis and so is not shown here.**

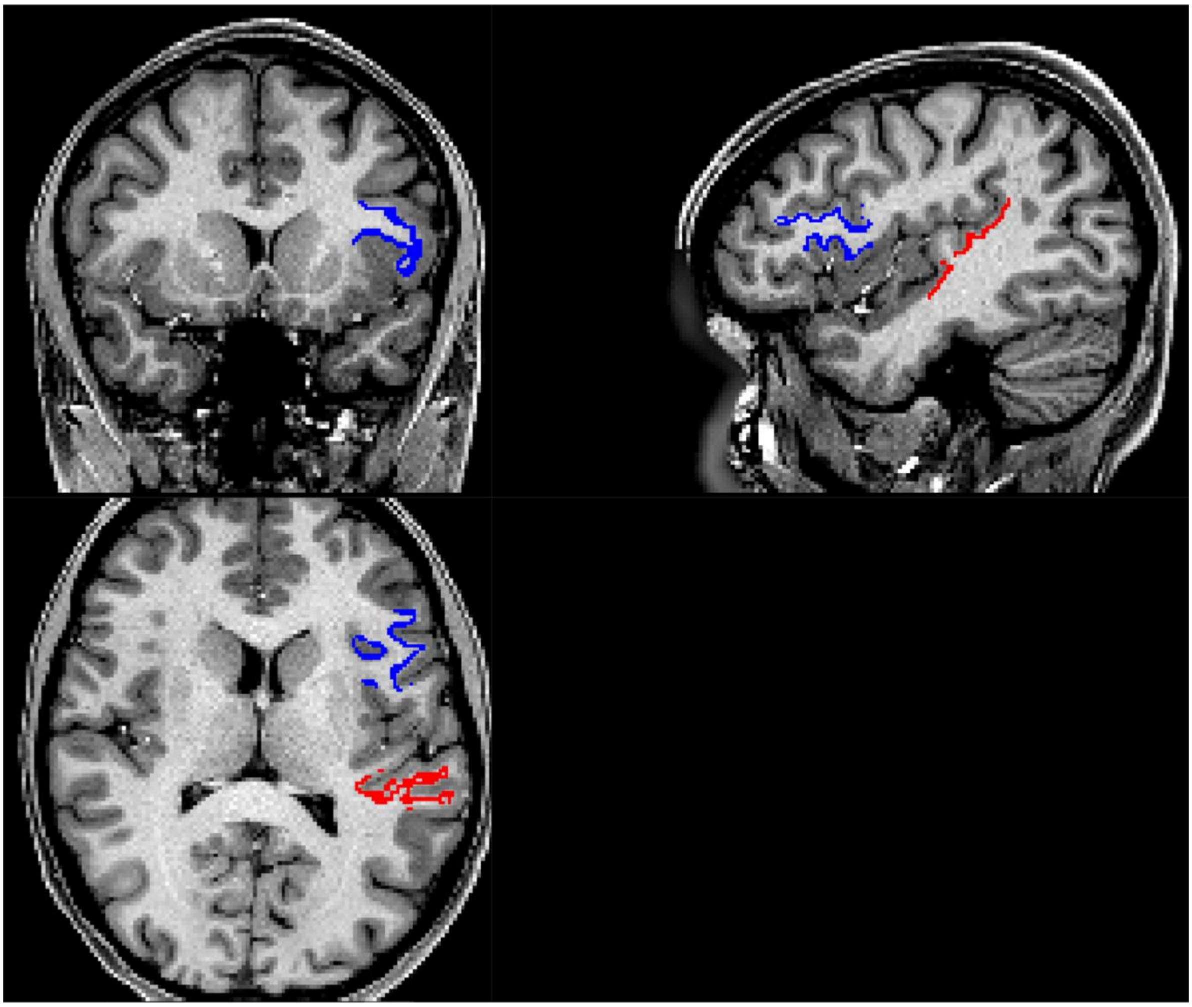

**Supplementary Figure 5. Example of a seed (blue) and superior temporal lobe inclusion mask (red) for generation of the arcuate fasciculus. Image is an unprocessed T1 MPRAGE.**

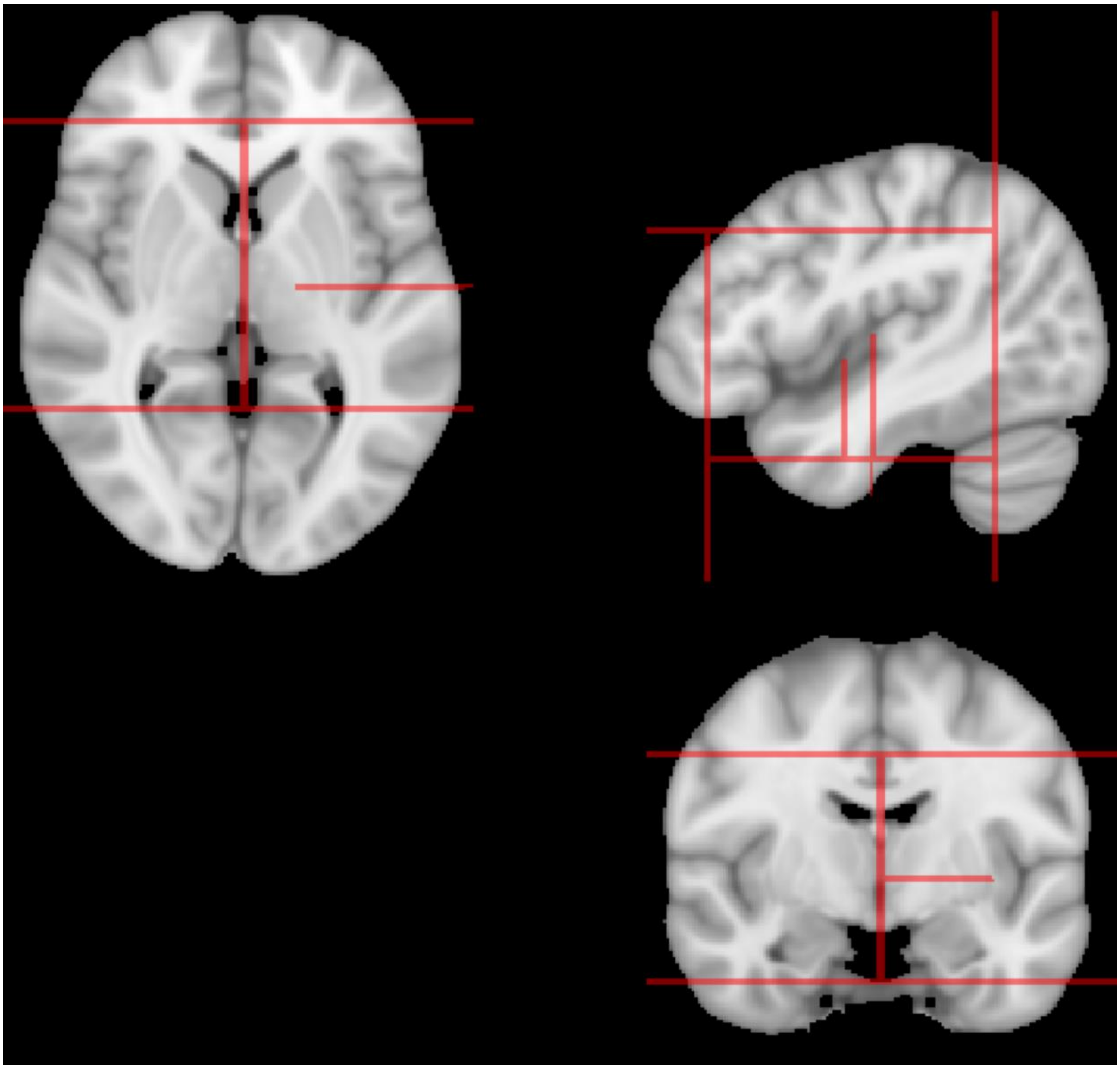

**Supplementary Figure 6. Exclusion mask for the arcuate fasciculus included several partial planes lying in each plane. The mask is not mirrored on the right side of the brain (left of image) because only the left tract was delineated. The exclusion mask predominantly ensured that the arcuate fasciculus, rather than other tracts, was delineated but also ensured that streamlines did not pass anteriorly through the temporal lobe and then superiorly into the frontal lobe.**

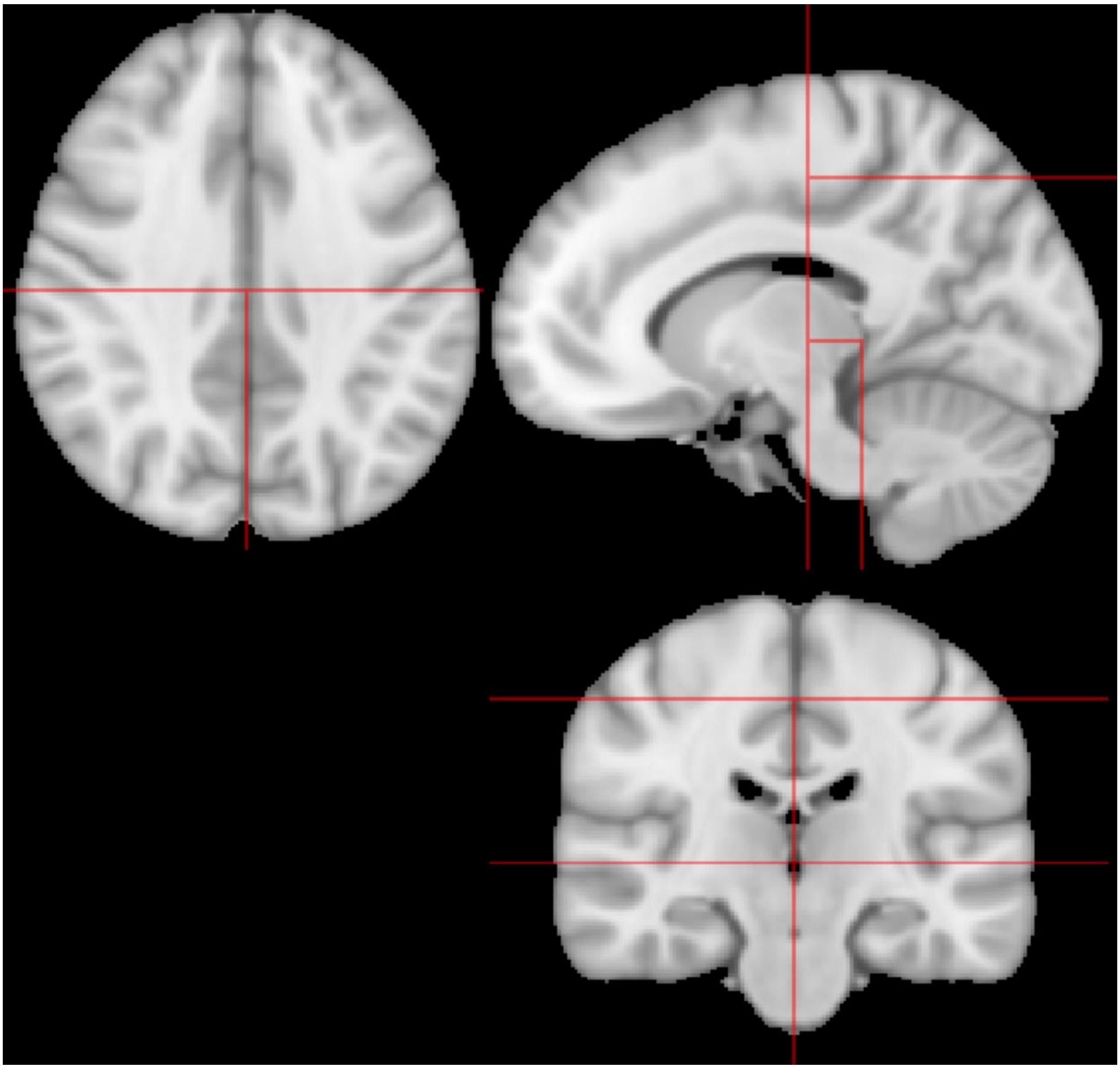

**Supplementary Figure 7. Exclusion mask for the forceps major included partial planes lying in the sagittal midline, axial plane at a location superior to the corpus callosum, and coronal plane at positions inferior to and anterior to the splenium of the corpus callosum. Masked voxels within the midbrain and brainstem prevented stray streamlines entering the brainstem and corpus callosum. The sagittal midline prevented streamlines crossing hemispheres through particularly thin aspects of the grey matter in visual cortex.**

### S1. Streamline Count vs Dice Coefficient

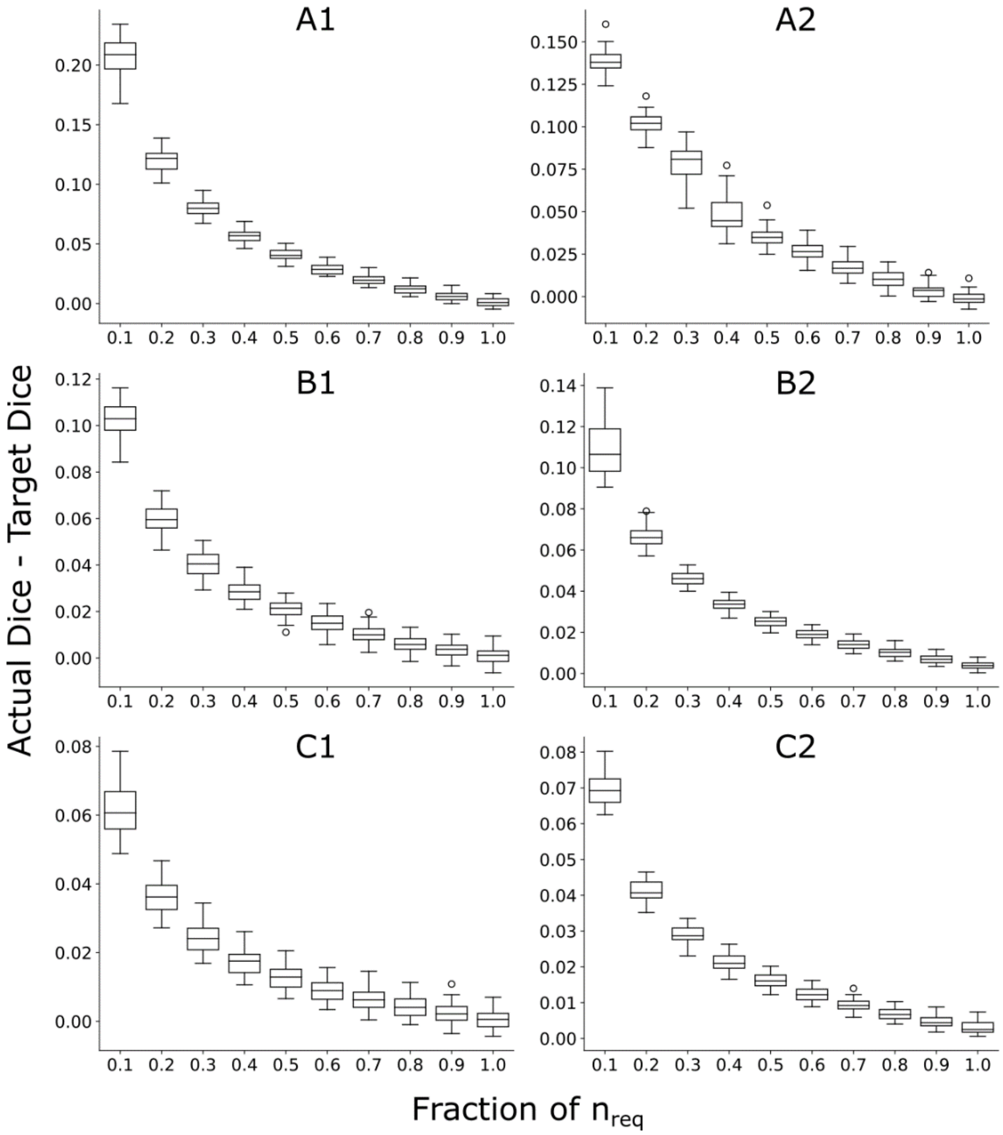


**Supplementary Figure 8. Relationship between streamline number, normalized to $n_{req}$, and tractography reliability of the arcuate fasciculus. The y-axis indicates the actual Dice coefficient minus the target Dice coefficient ($D_t$). Each participant contributed one datapoint to each box, in each plot. A1, A2, B1, B2, C1, and C2 refer to different stopping criteria, defined in Table 2. Error progressively reduced as more streamlines were added, reaching approximately zero for most stopping criteria when $n_{req}$ streamlines had been generated.**

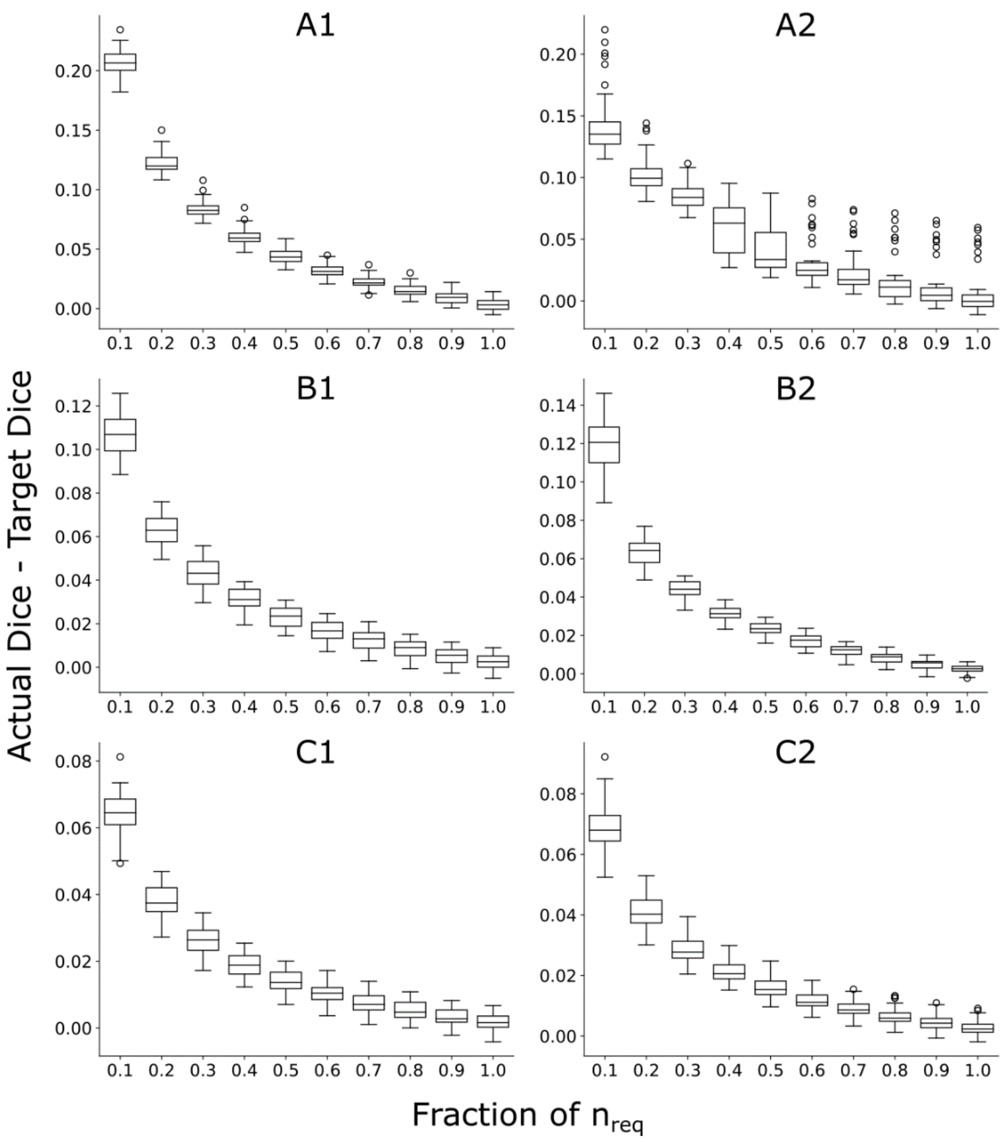


**Supplementary Figure 9. Relationship between streamline number, normalized to $n_{req}$, and tractography reliability of the forceps major. The y-axis indicates the actual Dice coefficient minus the target Dice coefficient ($D_t$). Each participant contributed one datapoint to each box, in each plot. A1, A2, B1, B2, C1, and C2 refer to different stopping criteria, defined in Table 2. Error progressively reduced as more streamlines were added, reaching approximately zero for most stopping criteria when $n_{req}$ streamlines had been generated.**

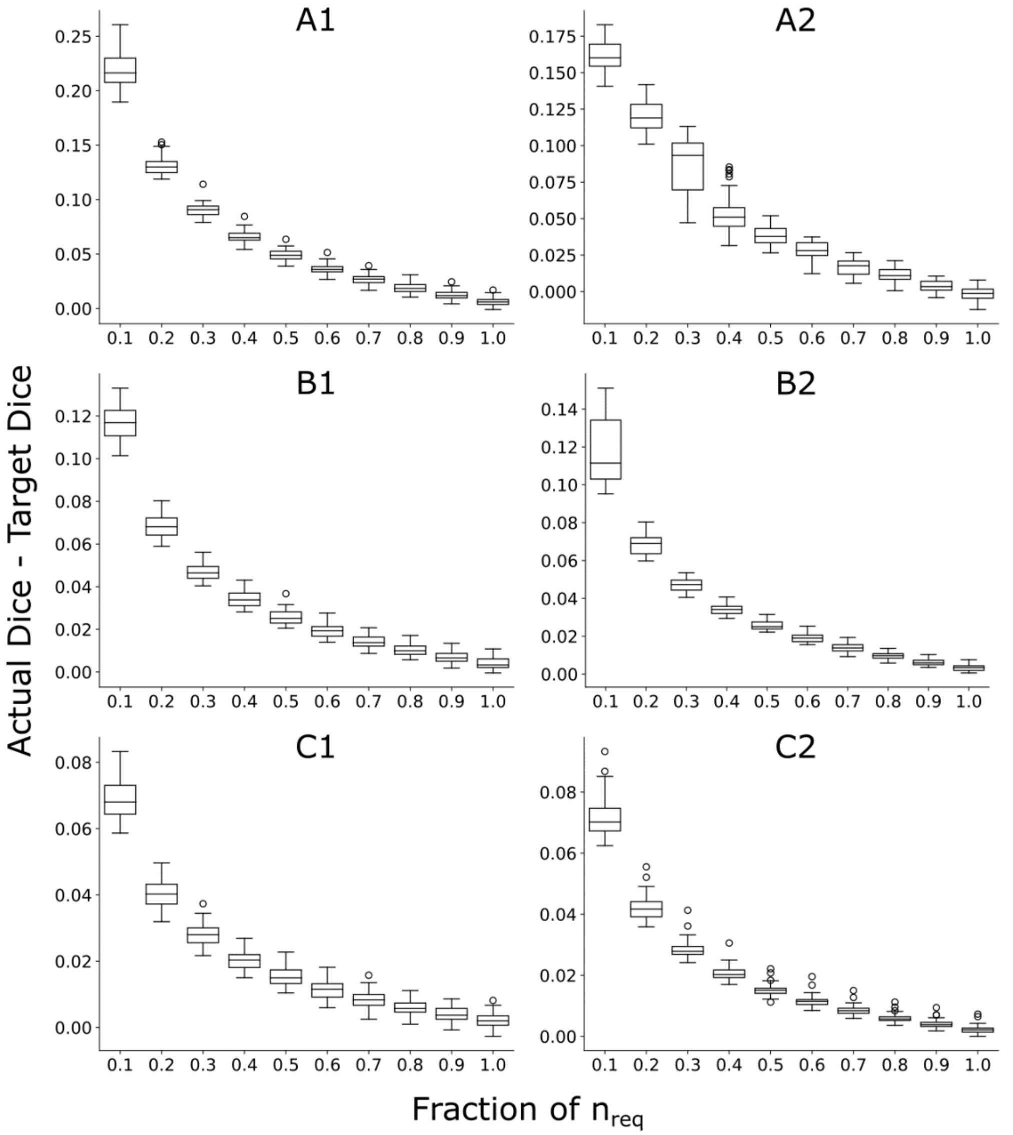


**Supplementary Figure 10. Relationship between streamline number, normalized to $n_{req}$, and tractography reliability of the corticospinal tract, at 1.25mm resolution. The y-axis indicates the actual Dice coefficient minus the target Dice coefficient ($D_t$). Each participant contributed one datapoint to each box, in each plot. A1, A2, B1, B2, C1, and C2 refer to different stopping criteria, defined in Table 2. Error progressively reduced as more streamlines were added, reaching approximately zero for most stopping criteria when $n_{req}$ streamlines had been generated.**

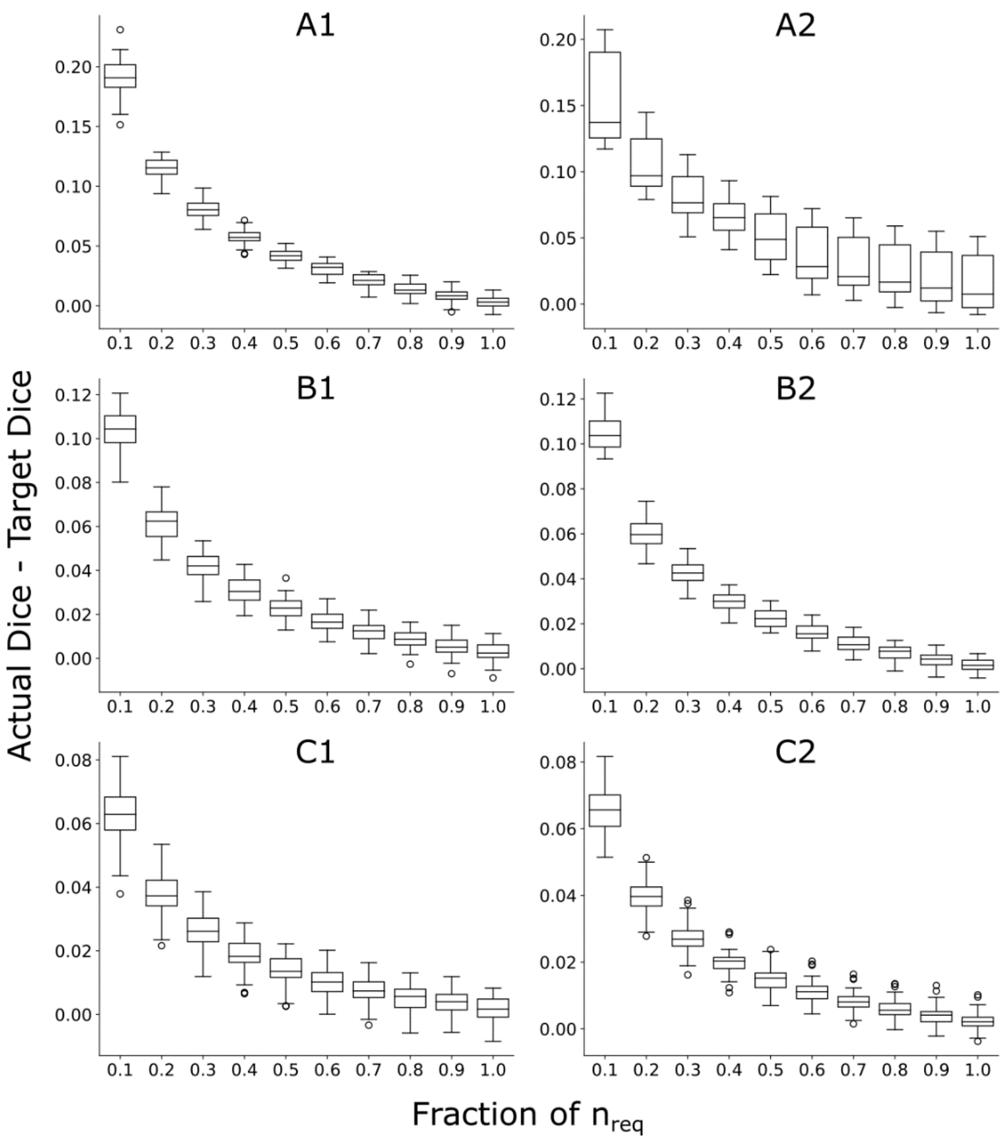


**Supplementary Figure 11. Relationship between streamline number, normalized to $n_{req}$, and tractography reliability of the corticospinal tract at 2mm resolution. The y-axis indicates the actual Dice coefficient minus the target Dice coefficient ($D_t$). Each participant contributed one datapoint to each box, in each plot. A1, A2, B1, B2, C1, and C2 refer to different stopping criteria, defined in Table 2. Error progressively reduced as more streamlines were added, reaching approximately zero for most stopping criteria when $n_{req}$ streamlines had been generated.**

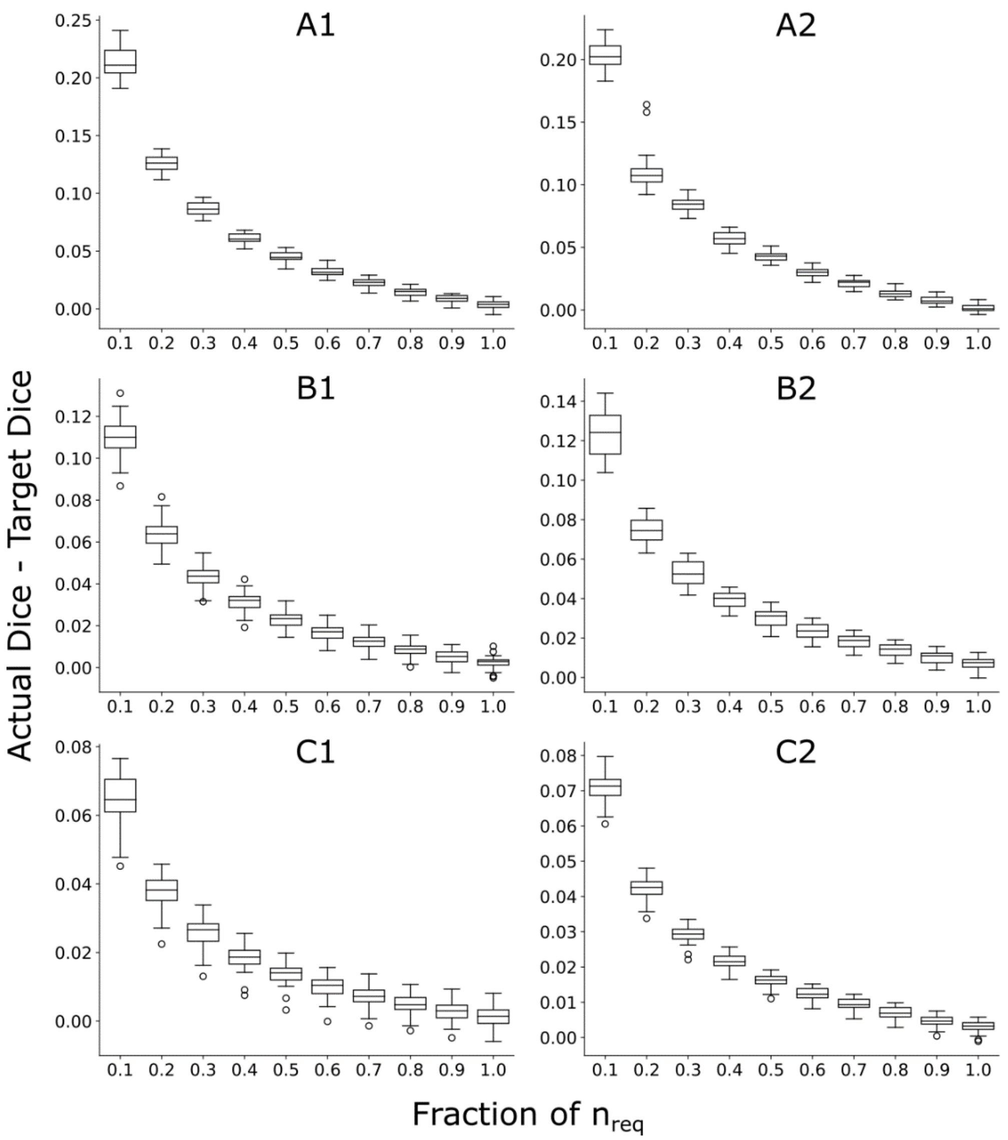


**Supplementary Figure 12. Relationship between streamline number, normalized to $n_{req}$, and tractography reliability of the corticospinal tract at 2mm resolution with a simulated single shell acquisition. The y-axis indicates the actual Dice coefficient minus the target Dice coefficient ($D_t$). Each participant contributed one datapoint to each box, in each plot. A1, A2, B1, B2, C1, and C2 refer to different stopping criteria, defined in Table 2. Error progressively reduced as more streamlines were added, reaching approximately zero for most stopping criteria when $n_{req}$ streamlines had been generated.**
